# Optimized directed evolution of *E. coli* leucyl-tRNA synthetase adds many noncanonical amino acids into the eukaryotic genetic code including ornithine and N^ε^-acetyl-methyllysine

**DOI:** 10.1101/2024.11.27.625662

**Authors:** Elise D. Ficaretta, Tarah J. Yared, Subrata Bhattacharjee, Lena A. Voss, Rachel L. Huang, Abhishek Chatterjee

## Abstract

Site-specific incorporation of noncanonical amino acids (ncAAs) into proteins in eukaryotes has predominantly relied on the pyrrolysyl-tRNA synthetase/tRNA pair. However, access to additional easily engineered pairs is crucial for expanding the structural diversity of the ncAA toolbox in eukaryotes. The *Escherichia coli*-derived leucyl-tRNA synthetase (EcLeuRS)/tRNA pair presents a particularly promising alternative. This pair has been engineered to charge a small yet structurally diverse group of ncAAs in eukaryotic cells. However, expanding the substrate scope of EcLeuRS has been difficult due to the suboptimal yeast-based directed evolution platform used for its engineering. In this study, we address this limitation by optimizing the yeast-based directed evolution platform for efficient selection of ncAA-selective EcLeuRS mutants. Using the optimized selection system, we demonstrate rapid isolation of many novel EcLeuRS mutants capable of incorporating various ncAAs in mammalian cells, including ornithine and N^ε^-acetyl-methyllysine, a recently discovered post-translational modification in mammalian cells.

## Introduction

The site-specific introduction of noncanonical amino acids (ncAA) into proteins in living cells using the genetic code expansion (GCE) technology is a powerful tool to probe and engineer protein function.^[1]^ GCE requires the use of an orthogonal (i.e., non-cross-reactive) aminoacyl-tRNA synthetase (aaRS)/tRNA pair, which co-translationally incorporates the desired ncAA in response to a repurposed nonsense codon.^[1]^ To ensure orthogonality, the ncAA-selective aaRS/tRNA pairs are typically imported into the host cell from a different domain of life. For example, GCE in bacteria has relied on pairs derived from archaea and eukaryotes, whereas the same in eukaryotic cells uses pairs derived from bacteria and archaea.

The GCE technology has been extended to eukaryotes, including in mammalian cells, where it has empowered new ways to probe the complex eukaryotic cell biology and engineer next-generation biotherapeutics.^[1-2]^ Several aaRS/tRNA pairs have been adapted for ncAA incorporation in eukaryotes, including the tyrosyl,^[3]^ leucyl,^[4]^ and tryptophanyl^[5]^ pairs derived from *E. coli*, the pyrrolysyl pairs from archaea,^[1c, 1f, 1g, 6]^ and chimeric pairs^[7]^ derived from bacterial and pyrrolysyl systems. Although hundreds of ncAAs have been introduced to the eukaryotic genetic code, the large majority of these rely on the pyrrolysyl pairs.^[1c, 1f, 6a]^ This impressive success stems from the intrinsic plasticity of PylRS and the ability to readily engineer it further using facile *E. coli*-based selection systems. However, access to other readily engineered pairs is needed to broaden the structural diversity of the ncAA toolbox in eukaryotic cells.

The *E. coli* leucyl-tRNA synthetase (EcLeuRS)/tRNA pair is a particularly attractive platform for GCE in eukaryotes. Since its development nearly 20 years ago, EcLeuRS has been engineered to genetically encode a small but structurally diverse group of ncAAs (Figure S1),^[1e, 3e, 4, 8]^ with sidechains that range from linear alkyl groups to complex aromatic ones. However, the size of the ncAA toolbox available through EcLeuRS is limited compared to PylRS. A key source of this limitation is the suboptimal nature of the yeast-based directed evolution platform used to engineer EcLeuRS.^[3b, 4]^ Two such yeast-based selections systems have been developed. One is a survival-based selection that uses a TAG-inactivated transcriptional activator GAL4 driving the expression of metabolic selection markers such as HIS3 and URA3.^[3b, 4, 9]^ The second is a FACS-based selection, involving the detection of a TAG-inactivated reporter displayed on the yeast cell surface.^[3e, 10]^ Survival-based strategies have been traditionally more popular for aaRS engineering, since these are less time- and resource-intensive, and have the potential to process larger libraries. Here, we show that the performance of the established survival-based selection system used for engineering EcLeuRS is compromised by leaky expression of the TAG-inactivated GAL4. By redesigning the selection scheme for improved stringency and performance, we enabled efficient enrichment of ncAA-selective EcLeuRS mutants after just a single round of positive and negative selection. This optimized selection system allowed us to identify many novel EcLeuRS mutants for charging 2-aminocaprylic acid (Cap **1**), a photocaged citrulline derivative (ONBC **2**), and N^ε^-acetyl-methyllysine (Kacme **3**), a recently discovered novel post-translational modification in mammalian cells.^[11]^ Furthermore, several of these new EcLeuRS mutants showed remarkable substrate polyspecificity, enabling incorporation of numerous additional ncAAs (Figure 1).

**Figure 1.**
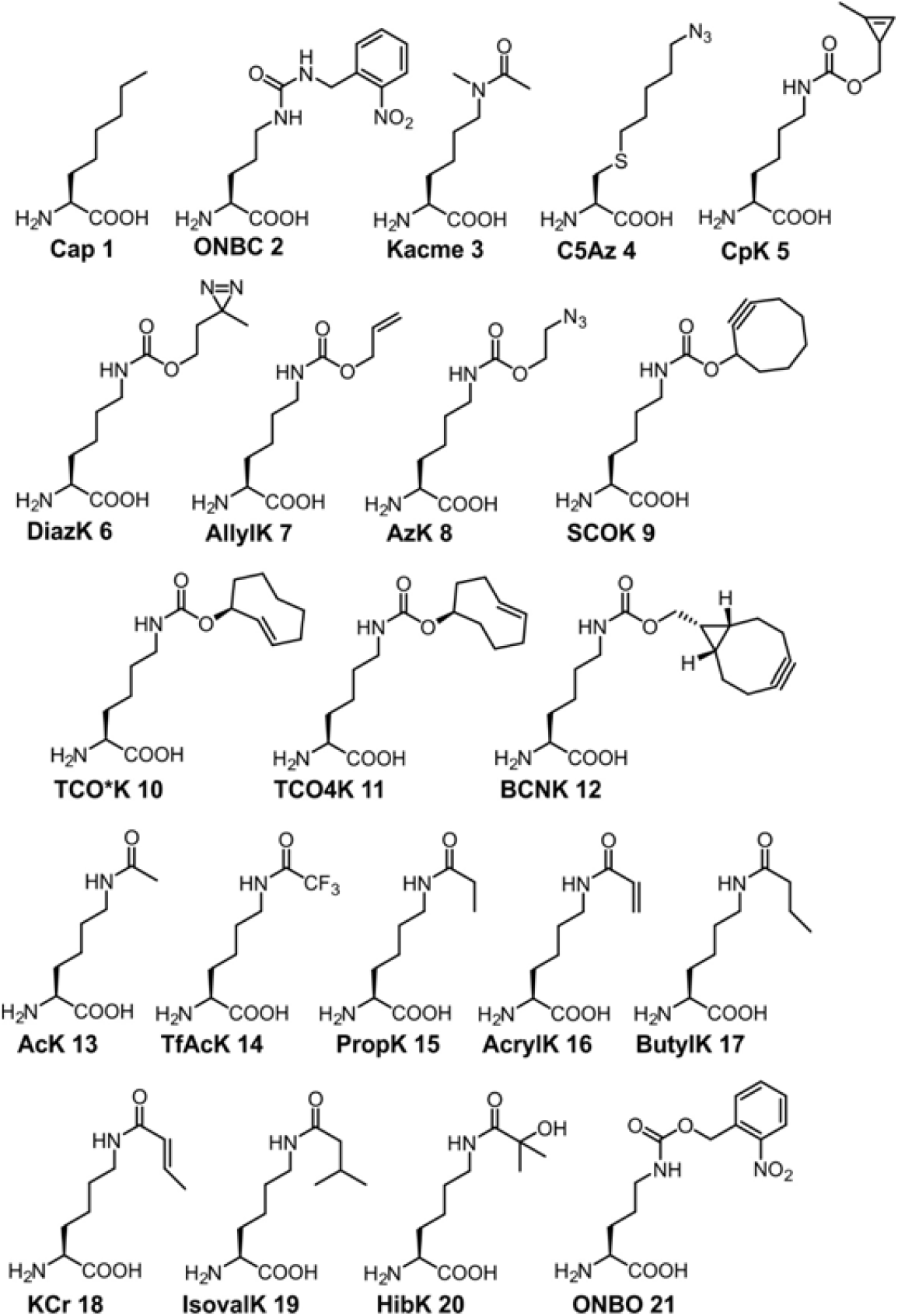
Structures of the ncAAs used in this work.

## Results and discussion

Because of its bacterial origin, the EcLeuRS/tRNA pair must be engineered in a eukaryotic host, where it would not cross-react with its endogenous counterparts. A yeast-based directed evolution system was developed for this purpose where the EcLeuRS/tRNA^EcLeu^_CUA_ pair suppresses TAG codons in the GAL4 transcription factor (GAL4-T44TAG-R110TAG; GAL4-2*), which subsequently drives the expression of metabolic reporters such as URA3 or HIS3 (Figure 2A).^[3b, 4, 9]^ Expression of these reporters allows the survival of yeast in media lacking uracil (Ura) and histidine (His), respectively, allowing selection of active EcLeuRS mutants. To deplete EcLeuRS mutants that remain active in the absence of the intended ncAA substrate, a negative selection is used where URA3 expression leads to the conversion of 5-fluoroorotic acid (5-FOA) to a toxic metabolite, resulting in cell death (Figure 2A). Several rounds of alternating positive and negative selection steps are typically used to enrich ncAA-selective EcLeuRS mutants before screening individual clones.

**Figure 2.**
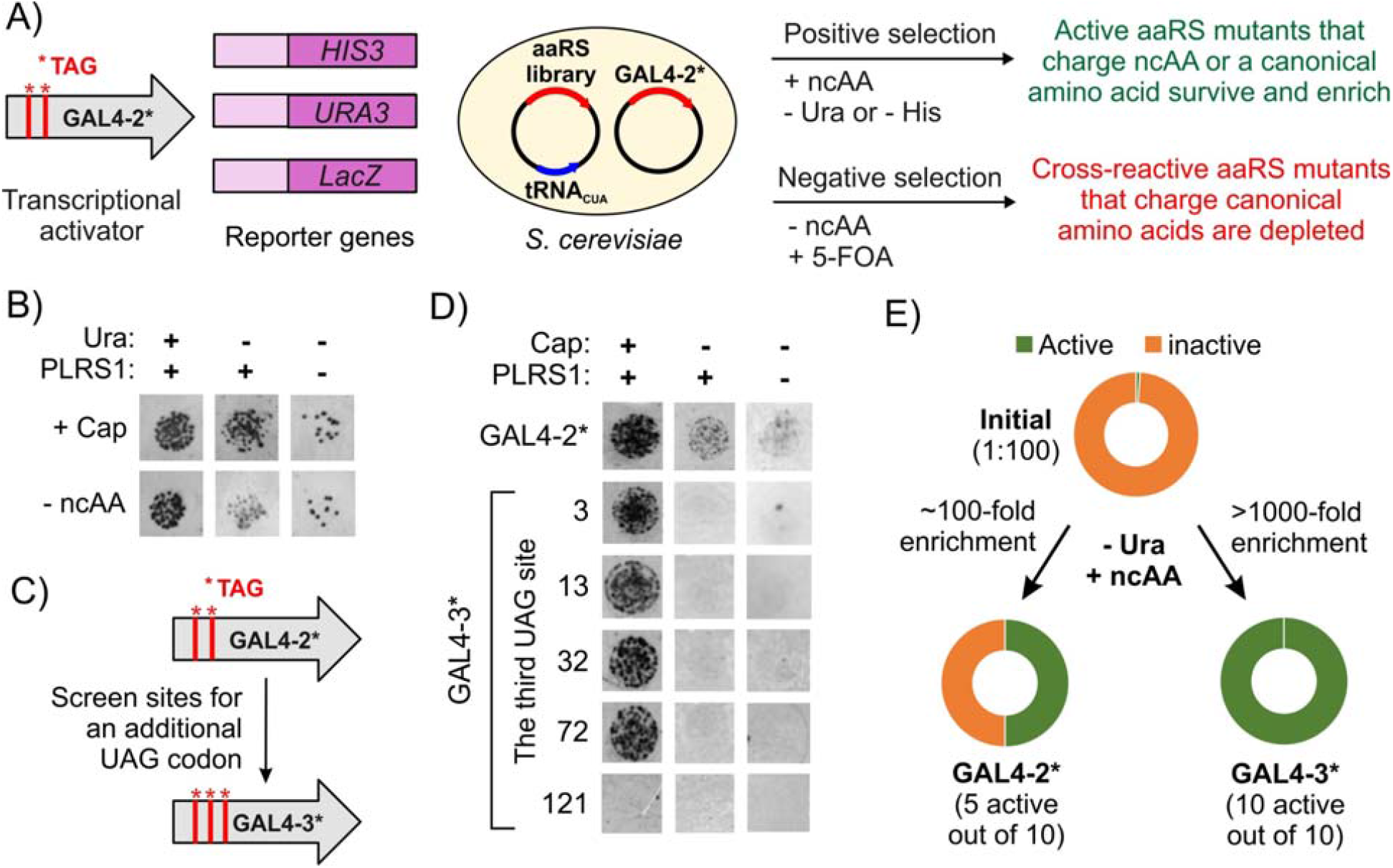
Improving the stringency of the yeast-based selection system. A) The survival-based selection scheme in yeast for engineering aaRSs. B) The established selection system using GAL4-2* is insufficiently stringent: In media lacking uracil, yeast cells expressing the PLRS1/tRNA^EcLeu^ CUA pair show robust growth in the presence of Cap (a substrate for PLRS1) as expected. However, weak but significant growth is also observed in the absence of Cap, as well as for yeast cells expressing only tRNA^EcLeu^ CUA but no EcLeuRS. C) Introduction of a 3^rd^ TAG codon in GAL4-2* was explored as a strategy to increase selection stringency and attenuate the leaky survival. D) Evaluating the performance of five GAL4-3* variants during positive selection. Four GAL4-3* variants enable robust growth only in the presence of the PLRS1/tRNA^EcLeu^_CUA_ pair and its substrate Cap. Unlike GAL4-2*, no growth is observed in the absence of Cap or PLRS1. E) Measuring the enrichment of active EcLeuRS mutants, from a defined mixture of active and inactive variants (in a 1:100 ratio), upon positive selection using the GAL4-2* or GAL4-3* selection system. Colony PCR of surviving clones after selection revealed a significantly higher degree enrichment of active EcLeuRS mutant using GAL4-3* (see Figure S2 for more details).

To benchmark the performance of this selection system, we used an established EcLeuRS mutant PLRS1,^[8h]^ which was previously shown to incorporate a variety of ncAAs including Cap. PLRS1 was co-expressed with its cognate suppressor tRNA_CUA_^EcLeu^ in yeast cells that also harbored the selection machinery (GAL4-2*, and GAL4-driven URA3, HIS3, and lacZ), and spot-plated on defined media under positive selection conditions (-Ura), in the presence or absence of Cap. Although the cells grew more robustly in the presence of Cap, substantial unexpected growth was also observed in its absence (Figure 2B). Furthermore, cells that did not express an active EcLeuRS (only tRNA^EcLeu^_CUA_ and the selection machinery) also showed residual growth under selective conditions, highlighting the insufficient stringency of the established selection system. A low level of cross-reactivity of tRNA^EcLeu^_CUA_ in yeast could be responsible for the unexpected survival in the absence of an EcLeuRS. We further assessed the performance of this selection system by measuring the enrichment level of an active EcLeuRS mutant within a defined mixture of active and inactive variants. Two plasmids encoding PLRS1 (active population) or no EcLeuRS (inactive population) were mixed in a 1:100 ratio and subjected it to this selection system, and surviving clones were individually screened by PCR to quantify the enrichment (Figure 2E, Figure S2). The selection was found to alter the ratio of active:inactive population from 1:100 to approximately 1:1, indicating a ∼100-fold enrichment. Such a modest level of enrichment may not be sufficient for facile identification of rare ncAA-selective clones from large mutant libraries, explaining the need for using many rounds of selections in the past.

We posited that the leaky expression of GAL4-2*, which underlies the insufficient stringency observed in our selection, may be mitigated by introducing an additional TAG stop codon at a permissive site (Figure 2C). To test this hypothesis, we introduced a third TAG codon in GAL4-2* at 5 different positions to create 5 GAL4-3* variants. These sites were selected based on its structure and previous research.^[9]^ When tested under positive selection conditions (-Ura), four of these mutants enabled robust survival of yeast cells in the presence of PLRS1 and its substrate Cap, but no growth was observed in the absence of either Cap or an active EcLeuRS, suggesting substantial improvement in stringency (Figure 2D). When tested under negative selection conditions (+Ura, +0.1% 5-FOA), one of these GAL4-3* mutants (GAL4-L3TAG-T44TAG-R110TAG) showed optimal survival pattern (Figure S3), and was selected for subsequent experiments. We evaluated the performance of the new GAL4-3*-based selection system using a mixed pool of active and inactive (in 1:100 ratio) EcLeuRS-encoding plasmids, as described above. All of the 10 randomly picked surviving clones were found to encode the active EcLeuRS, indicating at least a 1,000-fold enrichment (Figure 2E, Figure S2). These observations suggest that the GAL4-3*-based selection system should enable significantly more efficient directed evolution of EcLeuRS relative to the original GAL4-2TAG.

### Directed evolution of EcLeuRS using the new selection system

To test the efficacy of the more stringent GAL4-3*-based selection system for engineering EcLeuRS, we constructed a mutant library by randomizing five key residues in its active site: M40, L41, Y499, Y527, H527 (Figure 3A). Additionally, we introduced a T252A mutation in the editing domain, which has been shown to improve ncAA charging efficiency by this synthetase.^[26]^ After introducing the library into yeast cells harboring the GAL4-3*-based selection system, we first attempted to enrich EcLeuRS mutants that are selective for Cap as a proof-of-concept.

**Figure 3.**
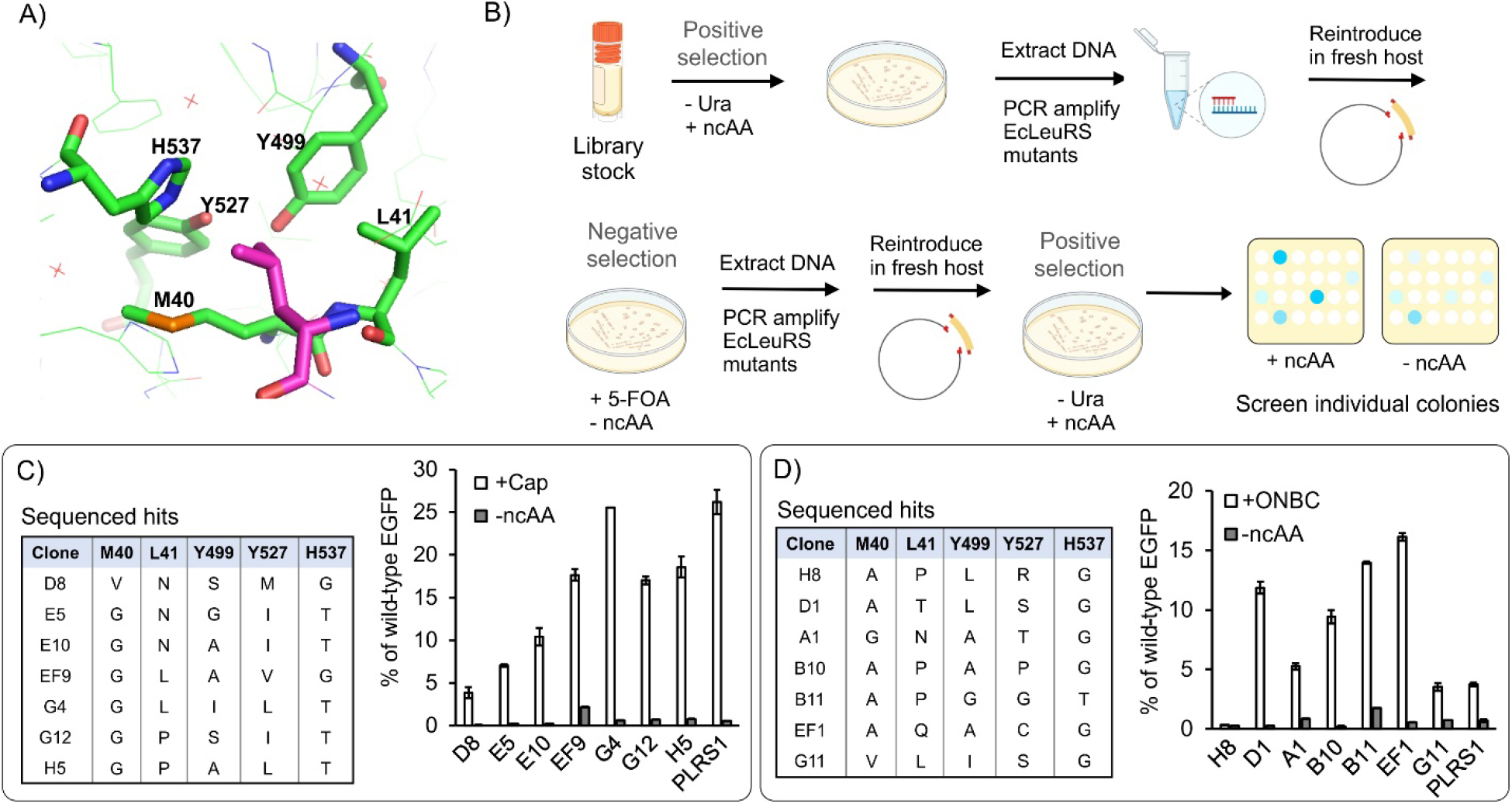
Engineering EcLeuRS using an optimized selection scheme. A) Structure of the EcLeuRS active site, highlighting the residues randomized in our library and the bound substrate (pink). B) A schematic of the optimized selection workflow using GAL4-3*, where surviving EcLeuRS mutant pool after a selection step is isolated and reintroduced into fresh host cells before the next round of selection. Robust enrichment of ncAA-selective mutants was observed after just three rounds (positive-negative-positive). C) Sequences of unique EcLeuRS variants that exhibited ncAA-dependent LacZ expression in yeast from the Cap selection, and the efficiency of Cap incorporation by these hits in HEK293T cells measured by the expression of the EGFP-39-TAG reporter. D) Sequences of unique EcLeuRS variants that exhibited ncAA-dependent LacZ expression in yeast after the ONBC selection, and the efficiency of ONBC incorporation by these hits in HEK293T cells measured by the expression of the EGFP-39-TAG reporter.

Initially, we followed previously described selection workflow, where yeast cells surviving one round of selection is directly used for the subsequent rounds following recovery.^[3b]^ However, we noted frequent appearance of a ‘cheating’ phenotype (clones that do not encode an ncAA-selective EcLeuRS, yet survive the selection), which gets progressively enriched in successive rounds of selection. We found that this issue could be addressed by isolating the EcLeuRS-encoding DNA from yeast cells after each selection step, and introducing them back into fresh host cells before proceeding to the next round of selection. Using this optimized selection scheme (Figure 3B), we observed highly efficient enrichment of ncAA-selective mutants after just three selective steps (positive-negative-positive). For example, when the library was selected for enriching Cap-selective mutants, numerous surviving clones after the 3^rd^ round of selection showed ncAA-dependent survival (Figure S4). Sequencing of ten such clones revealed seven distinct EcLeuRS mutants with similar active site architecture (Figure 3C). Next, these mutants were co-expressed in mammalian cells with tRNA^EcLeu^_CUA_ to test their ability to incorporate Cap into an EGFP-39-TAG reporter (Figure 3C). All seven mutants facilitated efficient Cap-dependent reporter expression, confirming that the selection was successful.

After confirming that our optimized selection system allows efficient directed evolution of EcLeuRS, we next focused on finding a mutant that efficiently charges ONBC **2**, a photocaged derivative of citrulline. Citrulline is an important PTM in eukaryotes, and its dysregulation is associated with numerous diseases such as rheumatoid arthritis.^[12]^ Protein arginine deiminases,^[13]^ enzymes responsible for introducing citrulline, are promiscuous, making it challenging to site-specifically incorporate this PTM into proteins. We previously found that the polyspecific EcLeuRS mutant PLRS1 fortuitously accepts ONBC, albeit with low efficiency.^[8d]^ Even though it enabled us to generate site-specifically citrullinated proteins for the first time, the low efficiency of this system has drastically limited its scope. To overcome this limitation, we sought to use our optimized selection system to develop an efficient ONBC-selective EcLeuRS mutant. Subjecting the aforementioned EcLeuRS library to the three-step selection as discussed above led to several clones that showed selective survival in the presence of ONBC (Figure S5). Sequencing of this clones the identification of seven unique mutants (Figure 3D). Six out of the seven mutants facilitated ONBC-dependent expression of the EGFP-39-TAG reporter in HEK293T cells (Figure 3D). Gratifyingly, five of these mutants were significantly more active than previously reported PLRS1, with the best two showing ∼4-fold improvement. Successful incorporation of ONBC was further confirmed by ESI-MS of the purified reporter protein (Figure S7). Discovery of this efficient citrulline incorporation system in mammalian cells will be useful to explore physiological consequences of numerous citrullination modifications that have been mapped in the mammalian proteome.^[14]^

### Genetically encoding the novel post-translational modification Kacme

Acetylation and methylation are established as two major post-translational modification (PTMs) of lysine in mammalian cells. Recently, Lu-Culligan *et al*. discovered that lysines can also be simultaneously acetylated and methylated to generate the novel PTM *N*^ε^-acetyl-*N*^ε^-methyllysine (Kacme) on histone 4 (H4), and it is associated with increased transcriptional initiation levels (Figure 4A).^[11]^ The physiological role of this novel PTM is poorly understood, and the ability site-specifically incorporate it into proteins will facilitate the studies toward elucidating its biology. We chemically synthesized Kacme and sought to use our selection system to identify an EcLeuRS that can charge this amino acid with good efficiency and selectivity. Subjecting our EcLeuRS library to three rounds of selection (positive, negative, positive) for Kacme incorporation, and sequencing the resulting clones that showed promising phenotype yielded 5 distinct mutants (Figure 4B). When tested in HEK293T cells using the EGFP-39-TAG reporter, all of the mutants exhibited Kacme-dependent reporter expression; clones H1 and EF9 in particular exhibited the highest activity (Figure 4C, 4D). Successful incorporation of Kacme was further verified by ESI-MS analysis of the reporter protein isolated using a C-terminal polyhistidine tag (Figure 4E, Figure S7). Finally, we used this technology to site-specifically introduce Kacme at two endogenous modification sites in H4 (K5 and K12). Plasmids encoding the H4-5-TAG and H4-12-TAG genes were individually transfected into in HEK293T cells with another plasmid harboring EF9-EcLeuRS and a recently engineered tRNA^EcLeu^_CUA_ variant LeuIGI (Figure 4F).^[15]^ Western blot analysis revealed successful expression of full-length H4 only in the presence of Kacme (Figure 4G).

**Figure 4.**
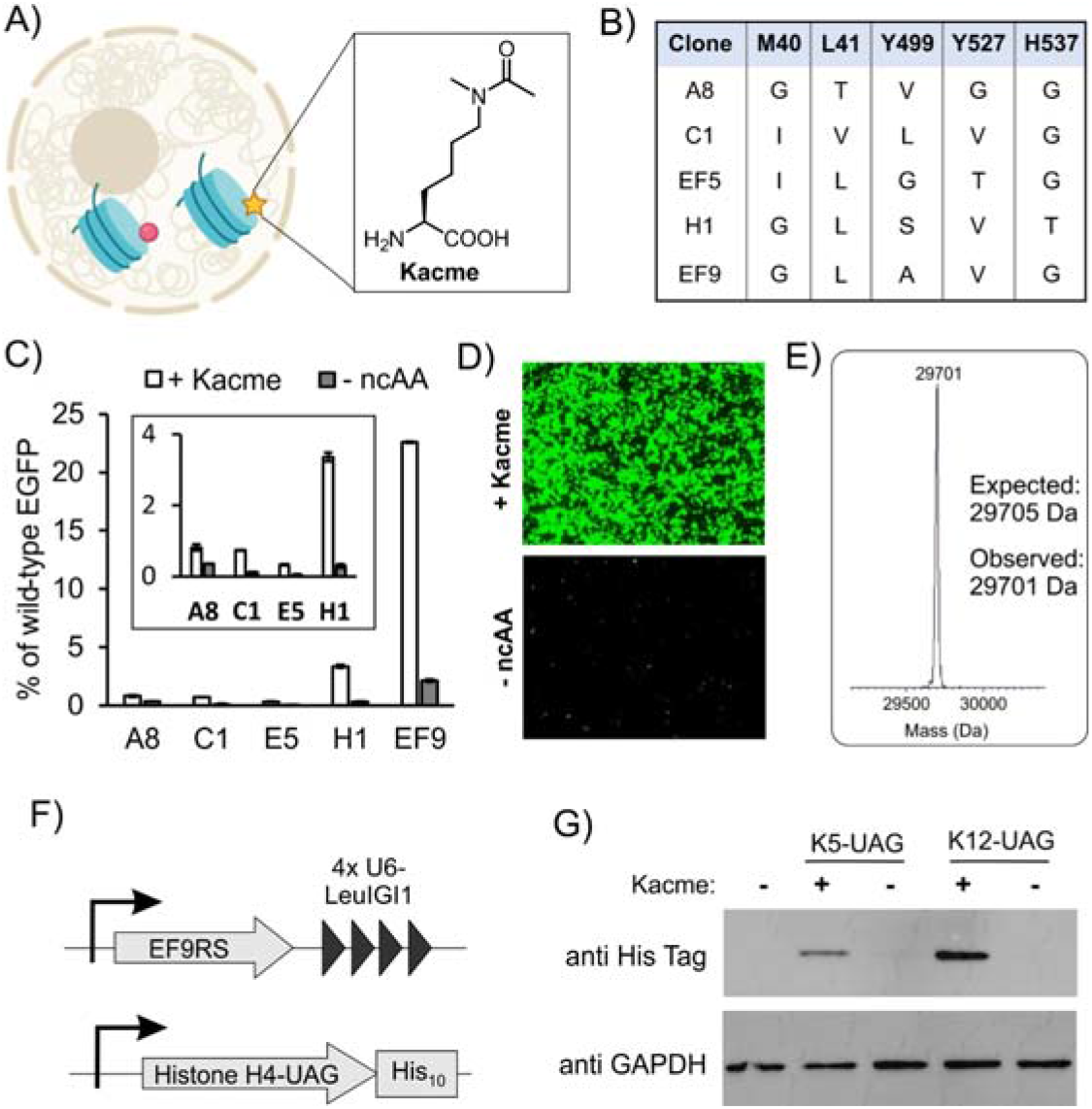
Genetically encoding Kacme in mammalian cells. A) Kacme has been recently identified as a unique histone H4 PTM. B) Sequences of unique EcLeuRS variants that exhibited ncAA-dependent LacZ expression in yeast after the Kacme selection. C) The efficiency of Kacme incorporation by these hits in HEK293T cells measured by the expression of the EGFP-39-TAG reporter. The results for hits A8, C1, E5, and H1 are magnified in inset. D) Fluorescence images of HEK293T cells co-transfected with EF9RS, tRNA^EcLeu^_CUA_, and EGFP-39-TAG in the presence and absence of Kacme. E) ESI-MS analysis of purified reporter protein corroborates successful Kacme incorporation. F) Constructs used for Kacme incorporation into histone H4. G) Western blot analysis shows Kacme-dependent expression of H4-5-TAG and H4-12-TAG, supporting successful incorporation of Kacme into two native sites in histone H4 where this PTM is found. GAPDH was used as a loading control.

### Identification of two super-polyspecific EcLeuRS mutants

It is known that certain aaRS mutants are polyspecific, meaning that they can accommodate a variety of structurally similar ncAAs.^[3c, 3d, 5a, 8h, 16]^ Such polyspecific aaRS mutants are valuable, because they enable facile incorporation numerous ncAAs without having to perform time- and resource-intensive directed evolution experiments. We suspected that the new EcLeuRS mutants reported here may exhibit significant substrate polyspecificity. Preliminary observations indicated that two particular EcLeuRS mutants, EF9 and B11, may be promising in this regard. EF9 was identified during both Cap and Kacme selections, whereas B11 appeared in the ONBC selection. We used a mammalian cell-based screen using the EGFP-39-TAG reporter to explore if EF9 and B11 can charge various linear aliphatic ncAAs. Both mutants exhibited remarkable substrate polyspecificity, allowing the incorporation of a wide range of ncAAs with diverse chemical functionalities (Figure 1, Figure 5A), including bioconjugation handles (e.g, CpK, SCOK, TCO*A, TCO4K, BCNK, C5Az), photoaffinity probes (e.g., DiazK), as well as a diverse family of PTMs such as AcK, TfAcK (hydrolysis-resistant AcK mimic), KCr, IsovalK, AcrylK, PropK, HibK, ButyrylK, etc. For a subset of these ncAAs, we used ESI-MS and SDS-PAGE analysis of the purified reporter protein to further confirm incorporation (Figure S6, S7).

**Figure 5.**
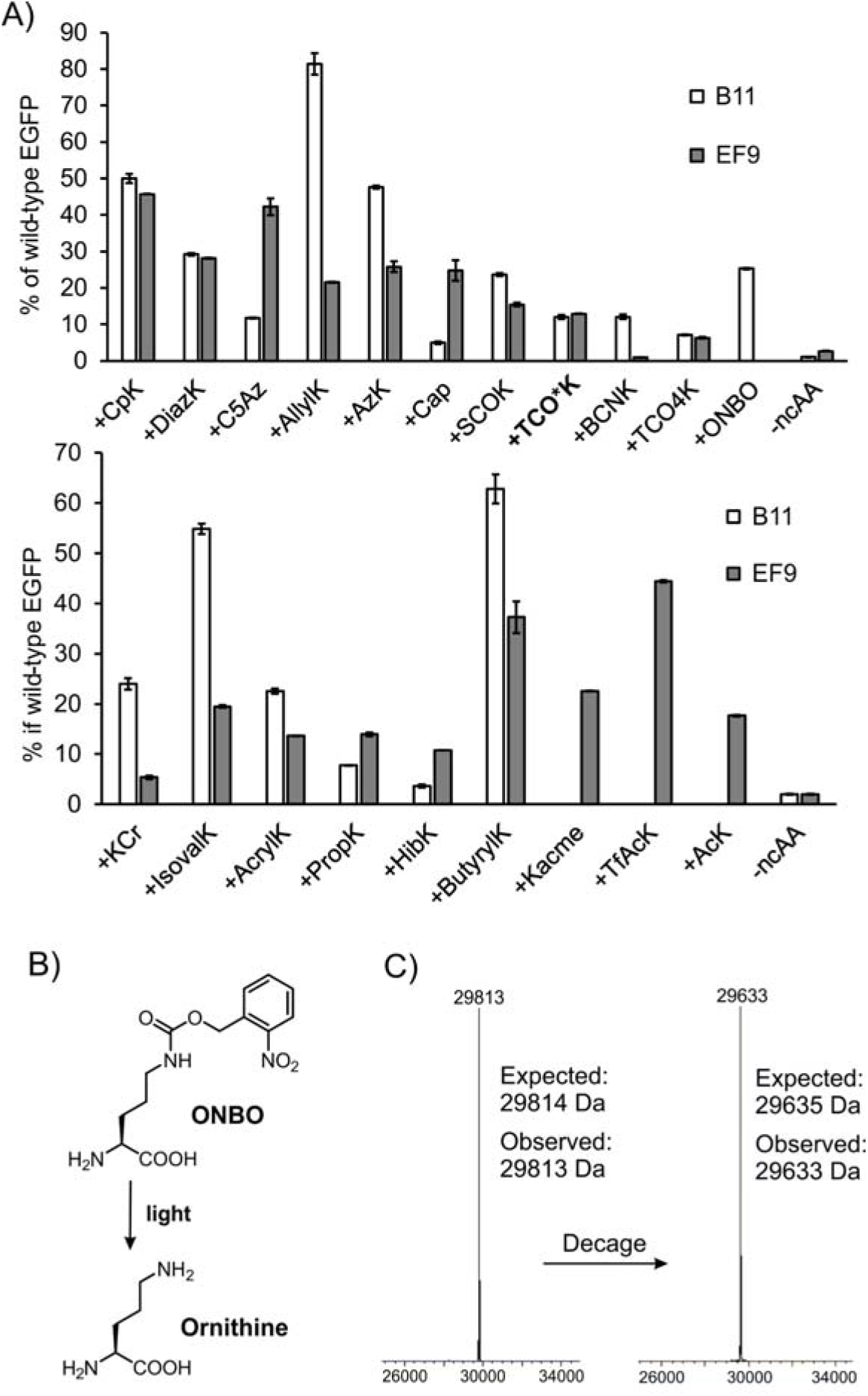
Remarkable substrate polyspecificity of two EcLeuRS mutants. A) Incorporation of various ncAAs using B11- and EF9-EcLeuRS in HEK293T cells measured by the expression of EGFP-39-TAG reporter using its characteristic fluorescence in cell-free extract. The lower panel includes various PTMs of lysine. Measurements were performed by co-transfecting pAAV-ITR-CMV-mCherry-1xU6-LeuIGI1 tRNA-plasmid, a pAcBac1-CMV-EGFP-39-TAG reporter plasmid, and a pIDT-CMV-EcLeuRS (B11 or EF9) plasmid into HEK293T cells, in the presence or absence of 10 mM (AcK), or 0.2 mM (for SCOK, TCO*A, TCO4K, BCNK, and ONBO), or 1 mM (all others) ncAA. Expression of EGFP-39-TAG was normalized relative to mCherry (encoded in the tRNA-expressing plasmid), and shown as a % of the wild-type EGFP reporter. Scheme (B) and ESI-MS analysis (C) of purified EGFP-39TAG-ONBO isolated from HEK293T cells using EcLeuRS hit B11 before (left) and after (right) photo-decaging to generate ornithine. Full spectra can be found in Figure S6.

The full substrate scope of the polyspecific EcLeuRS mutants is likely much larger; we only tested ncAAs that are either commercially available or were synthesized previously in the lab. Such ‘super-polyspecificity’ is reminiscent of some the most promiscuous PylRS mutants,^[16b]^ perhaps even exceeding them in some regard. For example, polyspecific PylRSs strongly prefer the presence of a carbonyl (e.g., an amide or carbamate) at the ^ε^-position within its substrates, while our polyspecific EcLeuRSs are less dependent on such specific features. Furthermore, we have previously shown that the EcLeu and the pyrrolysyl pairs are mutually orthogonal, allowing their simultaneous use to incorporate two distinct ncAAs into one polypeptide.^[8g, 8h, 17]^ The scope of such dual-ncAA applications is significantly expanded by adding many new ncAAs into the toolbox of the EcLeu pair.

Ornithine is an abundant non-proteinogenic amino acid, which can be also be generated post-translationally on polypeptides from an arginine residue.^[18]^ Site-specific incorporation of this shorter lysine homolog into proteins offers unique opportunities as a biochemical probe, as well as for protein engineering. Numerous bioactive peptide-derived natural products, including Gramicidin S,^[19]^ Tyrocidines,^[20]^ Daptomycin,^[21]^ Landornamide,^[22]^ etc., contain this unique amino acid, highlighting its rich utilization by nature for creating novel biological functions. However, the chemical instability of the putative ornithine-tRNA complex, which can decompose to generate a δ-lactam through a facile six-membered cyclic intermediate,^[23]^ makes its direct incorporation into proteins challenging. We envisioned that a photocaged derivative of ornithine, ONBO **21**, may overcome this challenge (Figure 5B). Upon its co-translational incorporation into proteins, ONBO can be efficiently converted to ornithine using light. Additionally, we suspected that ONBO, which is structurally similar to ONBC, may be readily accepted by the polyspecific B11-EcLeuRS. We chemically synthesized ONBO and tested this hypothesis to show that indeed B11-EcLeuRS efficiently incorporates ONBO into EGFP-39-TAG (Figure 5A). We confirmed the incorporation of ONBO by ESI-MS analysis of the purified reporter protein, and further demonstrated seamless removal of the photocage upon irradiation to generate the ornithine residue at the intended site (Figure 5C).

## Conclusions

The EcLeuRS/tRNA^EcLeu^ pair represents a highly promising platform for introducing structurally diverse ncAAs into the eukaryotic genetic code. However, its scope has been restricted by the suboptimal nature of the selection system needed to engineer it. Here, we have systematically identified and overcome these bottlenecks to create an optimized selection system that enables rapid enrichment of engineered EcLeuRS mutants with diverse substrate specificities. We demonstrate the utility of this platform by performing three selection experiments using Cap, ONBC, and Kacme, and retrieving several ncAA-selective EcLeuRS mutants in each case. Some of these mutants exhibit remarkable substrate polyspecificity, enabling the incorporation numerous ncAAs in mammalian cells, with the potential to accept many more substrates with similar architecture. Access to these polyspecific EcLeuRS mutants, and the ability to engineer the EcLeuRS/tRNA^EcLeu^ pair further using our facile directed evolution platform will significantly expand the scope of the GCE toolbox in eukaryotes.

## Acknowledgements

This work was supported by NIH (R35GM136437). The MaV203 yeast strain and pGADGAL4-2TAG and pESC-EcLeuRS-1xSUP4 plasmids were kind gifts from Prof. Peter G. Schultz

The authors have cited additional references within the Supporting Information.^[24]^

