## Supporting information for "Optimized directed evolution of *E. coli* leucyl-tRNA synthetase adds many noncanonical amino acids into the eukaryotic genetic code including ornithine and N^ε^-acetyl-methyllysine"

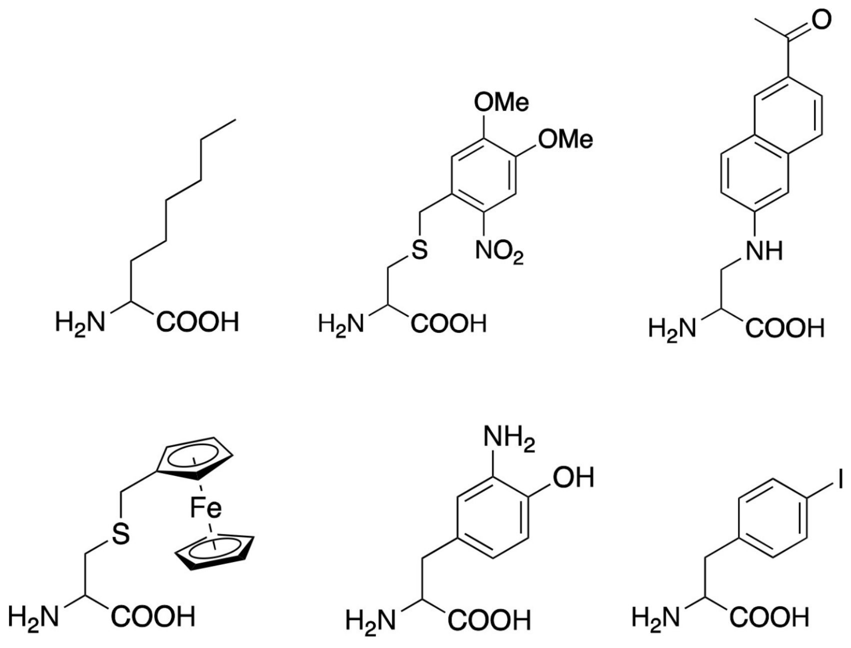


**Figure S1**. Examples of the structurally diverse ncAAs genetically encoded in eukaryotes using the *E. coli* leucyl pair.

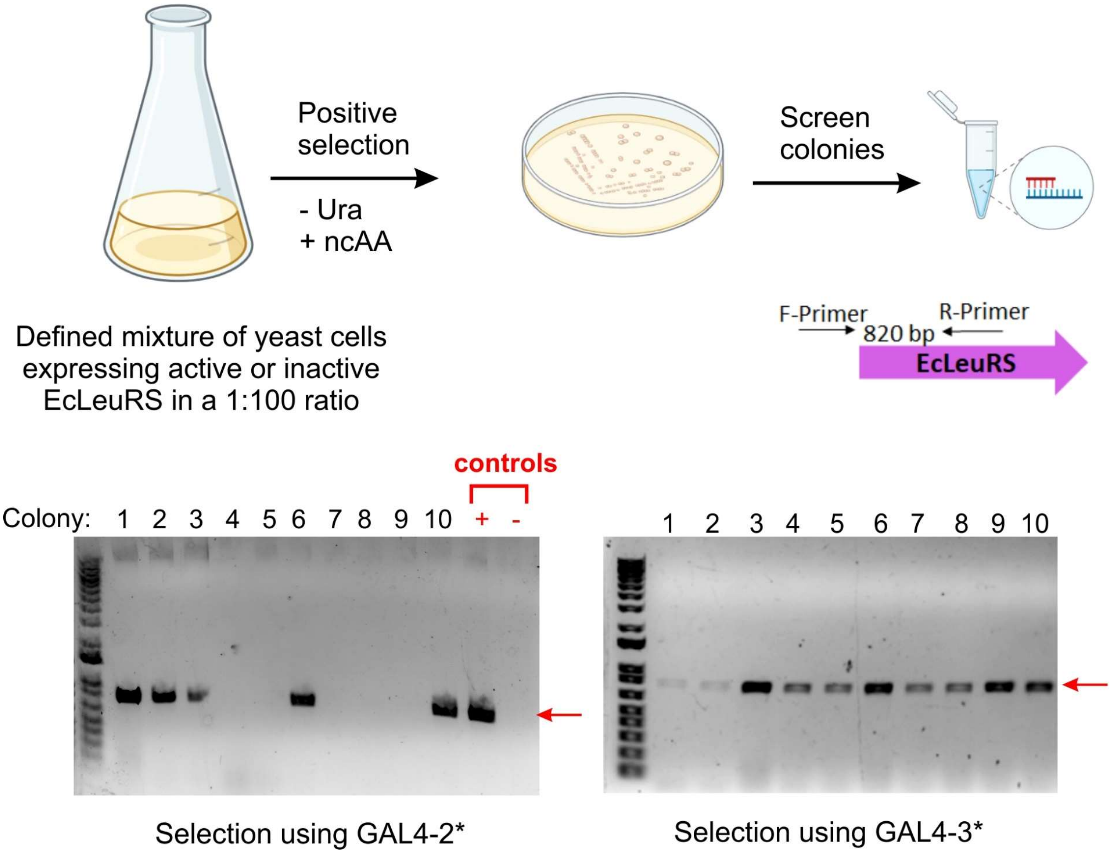


**Figure S2**. Measuring the enrichment of an active EcLeuRS mutant (PLRS1), when a defined mixture of yeast cells harboring either a PLRS1-encoding plasmid (active) or an empty (no EcLeuRS) plasmid (inactive) were subjected to the positive selection using a GAL4-2* or GAL4-3* system. The panel above shows the experimental scheme and the panel below shows the PCR screen of 10 randomly picked surviving colonies for the presence of active PLRS1. For GAL4-2* selection (left), 5 out of the 10 colonies that were screened yielded the expected band from PLRS1. For GAL4-3* selection (right), all of the 10 colonies that were screened yielded the expected band from PLRS1.

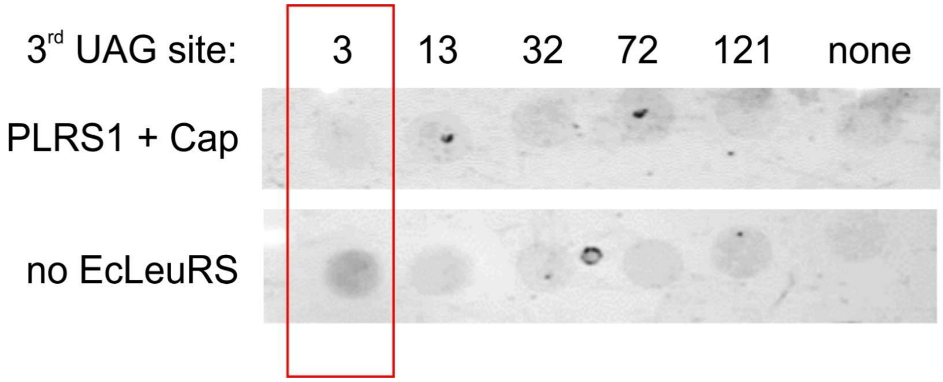


**Figure S3**. Evaluating the performance of the 5 GAL4-3* constructs under negative selection conditions (+ 5-FOA). The highlighted GAL4-3* construct exhibits the desired phenotype: No growth in the presence of an active EcLeuRS, but robust survival in the absence of one.

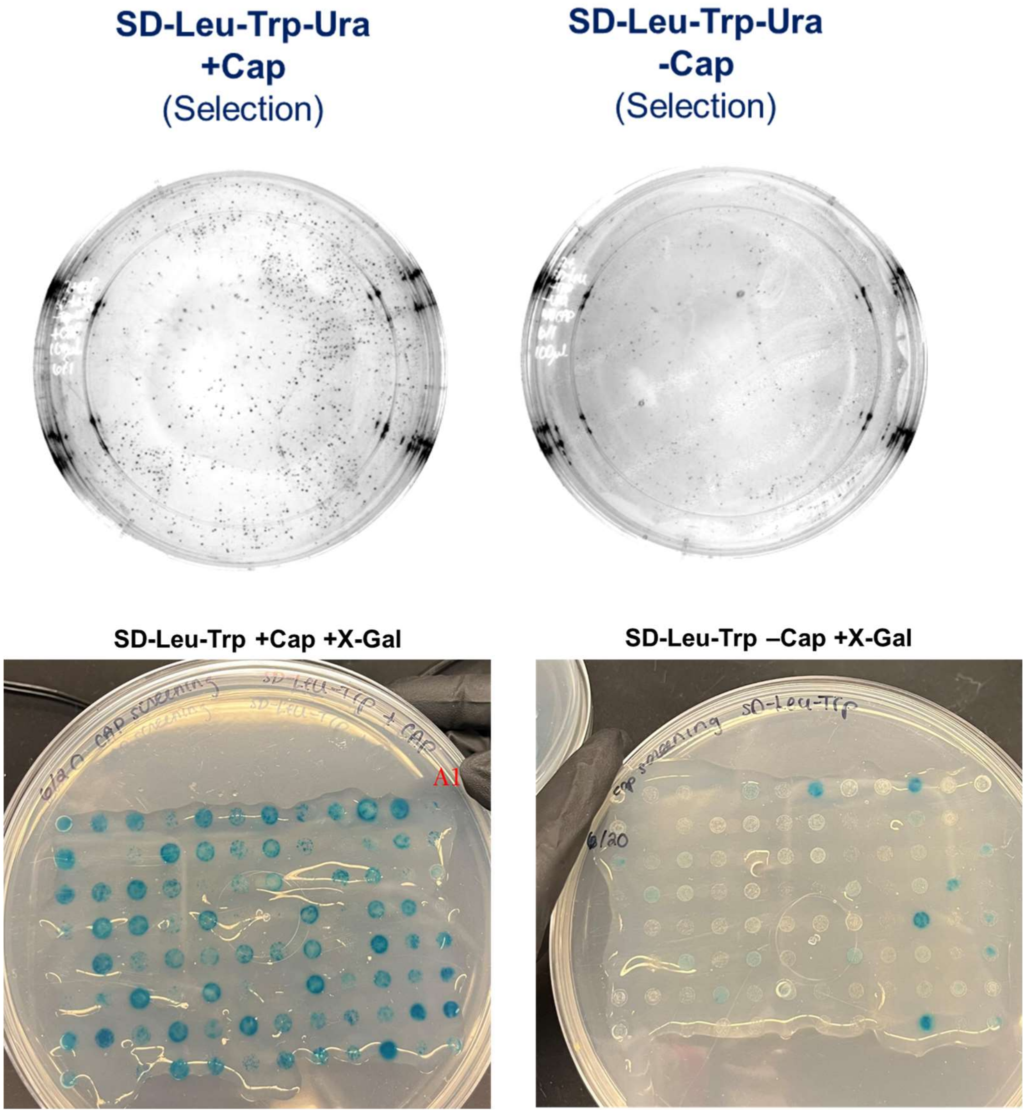


**Figure S4**. Successful selection of our EcLeuRS mutant library for charging Cap. Top panel: Images of the plates at the final round of positive selection show significant Cap-dependent growth. Bottom panel: Screening 96 individual colonies from the +Cap plate to look for Cap-dependent LacZ expression (visualized using the blue/white X-gal assay).

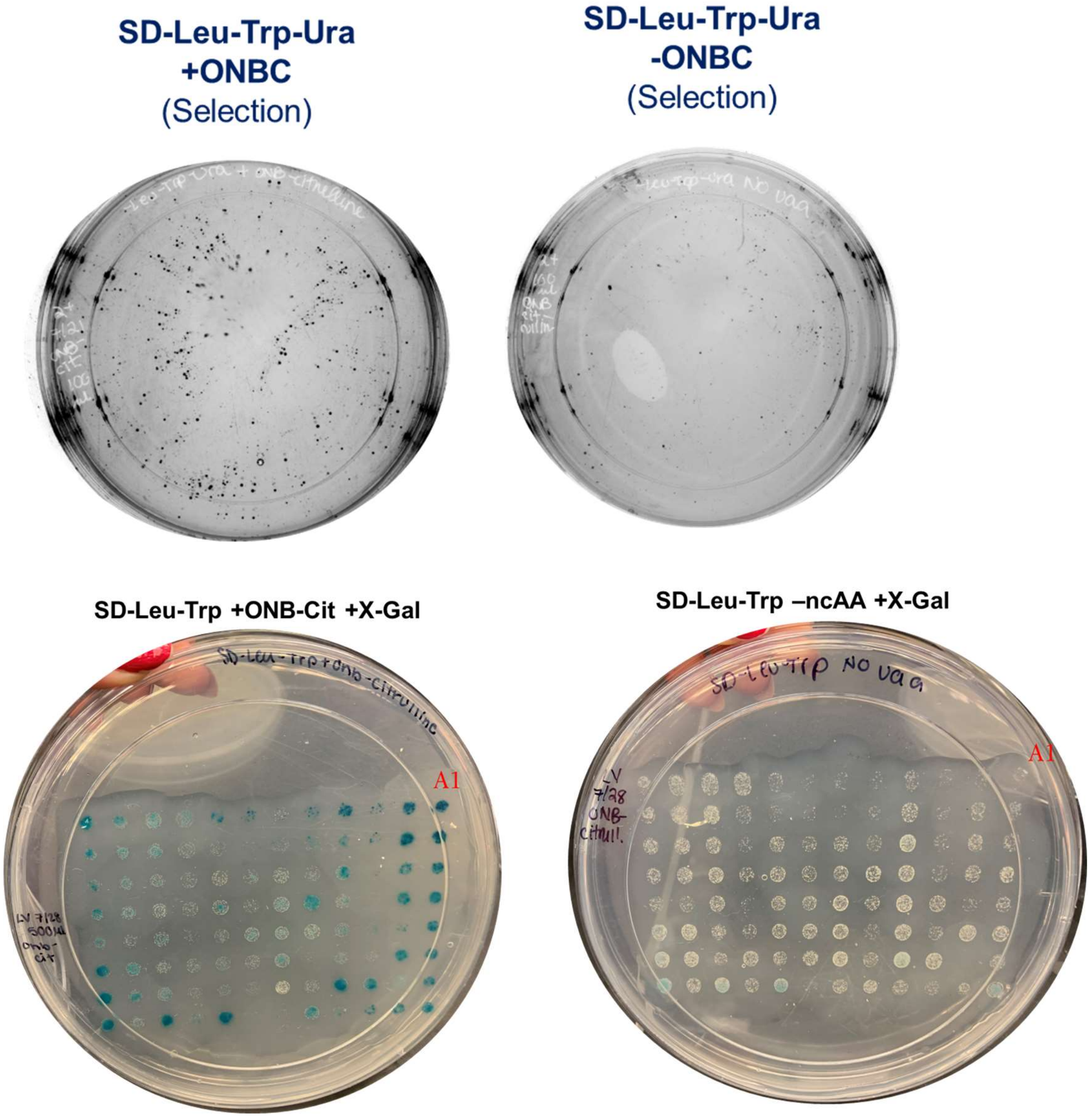


**Figure S5**. Successful selection of our EcLeuRS mutant library for charging ONBC. Top panel: Images of the plates at the final round of positive selection show significant ONBC-dependent growth. Bottom panel: Screening 96 individual colonies from the +ONBC plate to look for ONBC-dependent LacZ expression (visualized using the blue/white X-gal assay).

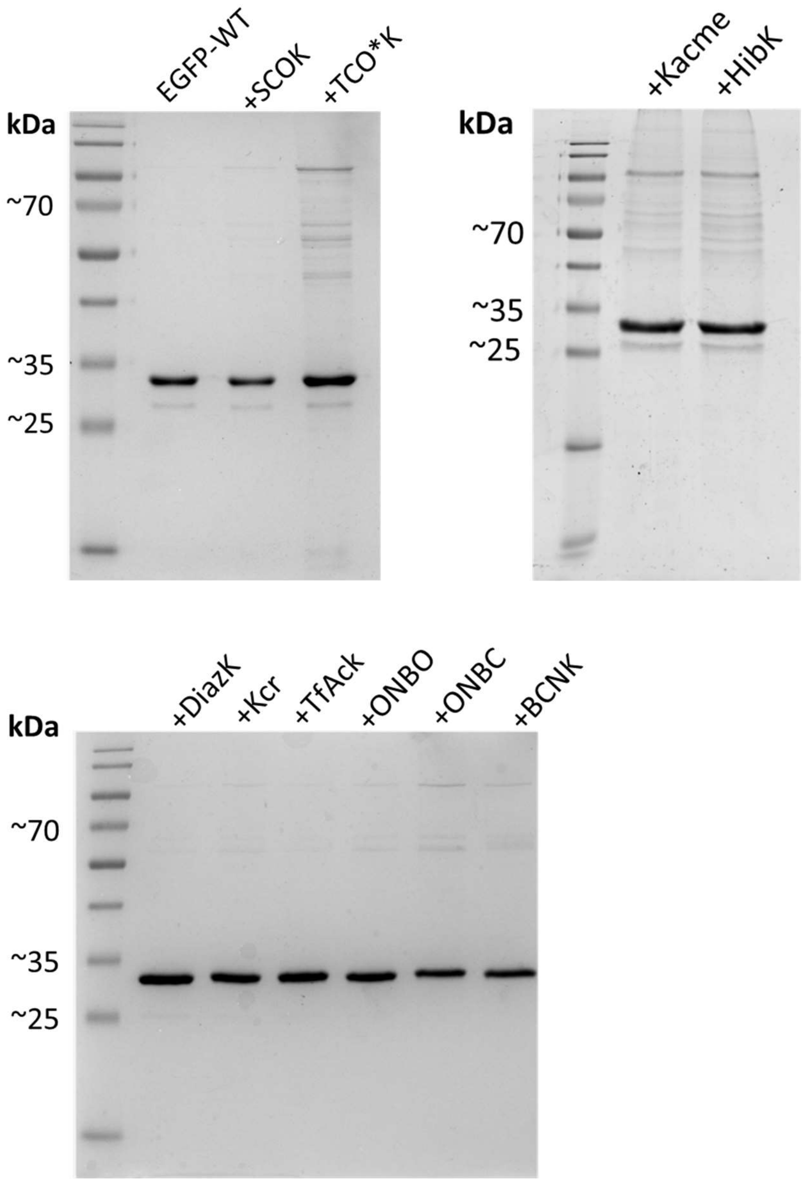


**Figure S6**. SDS-PAGE analysis of purified EGFP-39-TAG reporters incorporating various ncAAs purified from HEK293T cells. The expected size of the reporter is ~29 kDa.

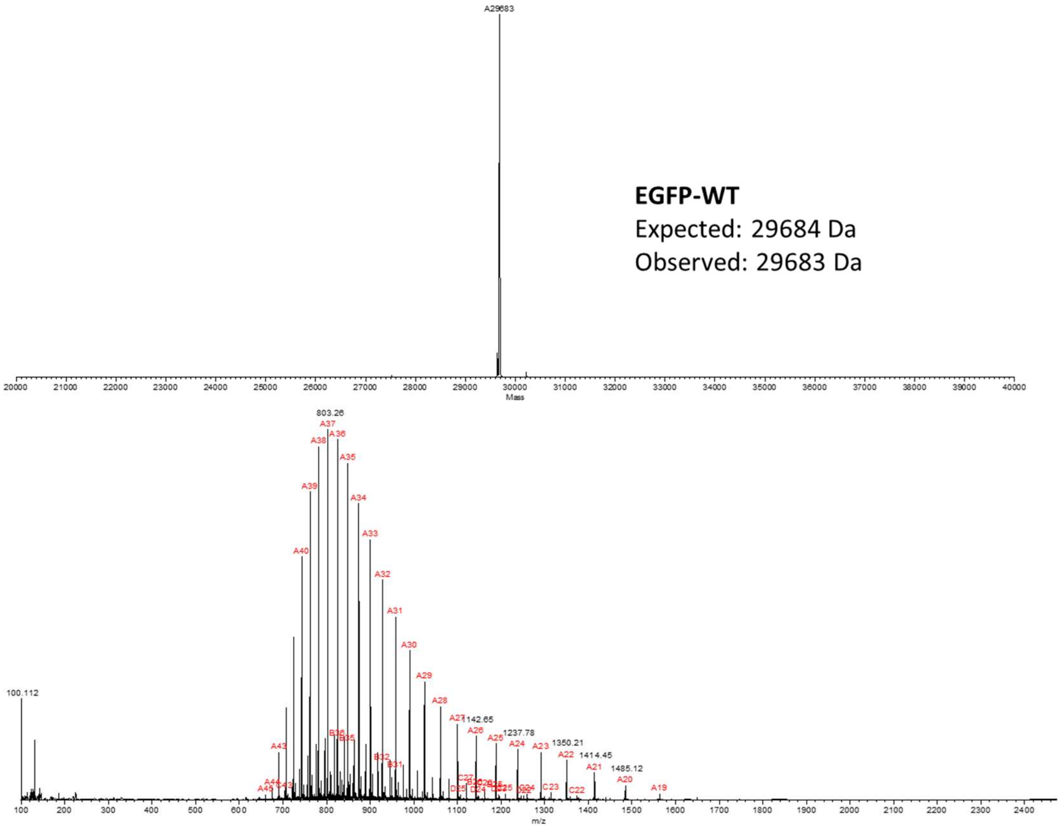


**EGFP-WT**
Expected: 29684 Da
Observed: 29683 Da

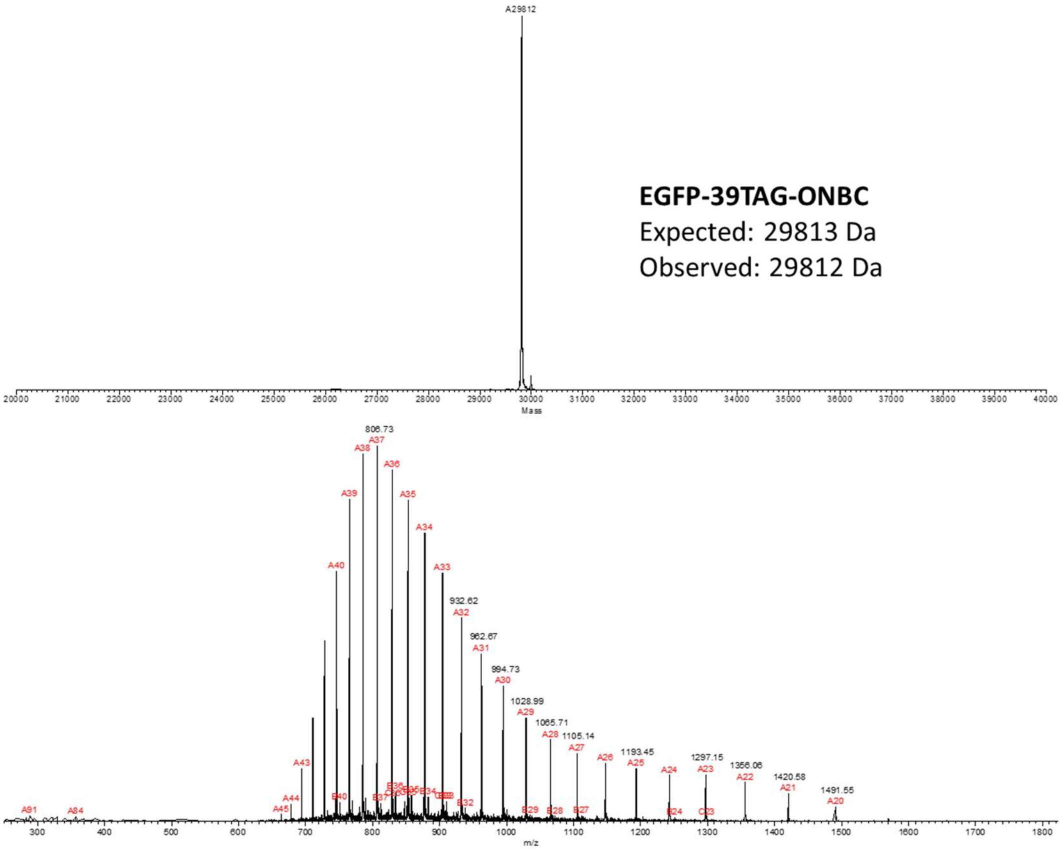


**EGFP-39TAG-ONBC**
Expected: 29813 Da
Observed: 29812 Da

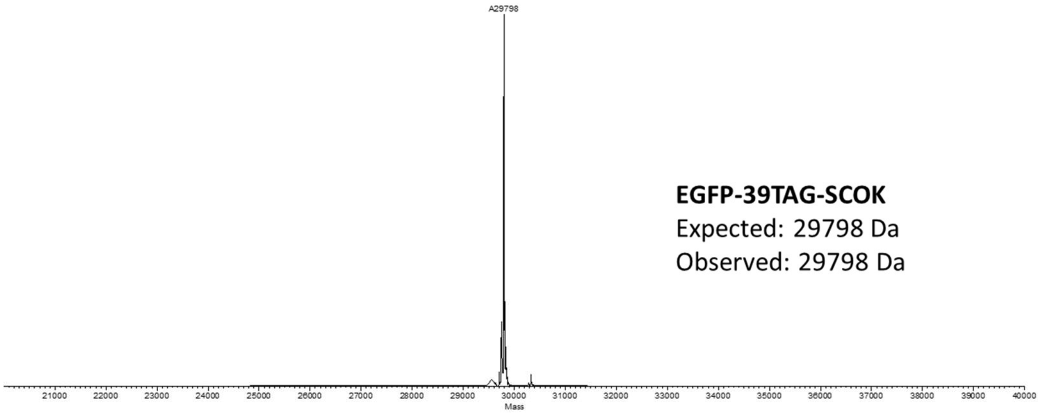
A29798
EGFP-39TAG-SCOK
Expected: 29798 Da
Observed: 29798 Da
Mass

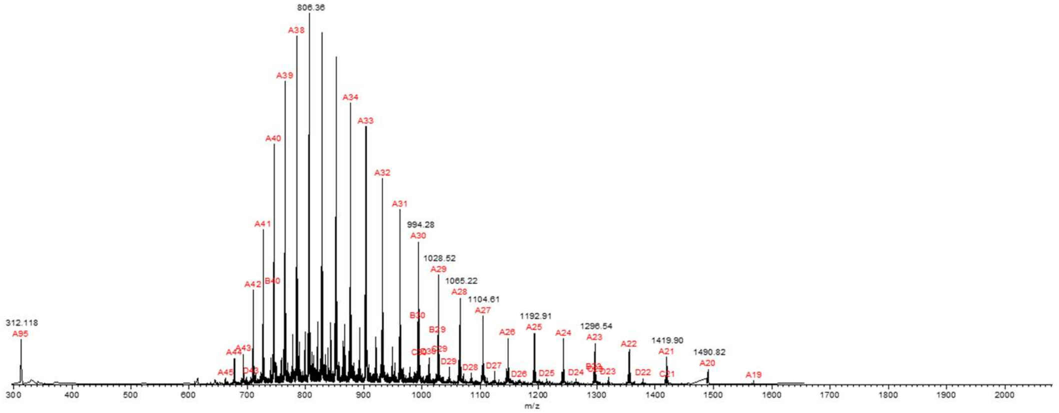
806.36
994.28
1028.52
1065.22
1104.61
1192.91
1296.54
1419.90
1490.82
312.118
m/z

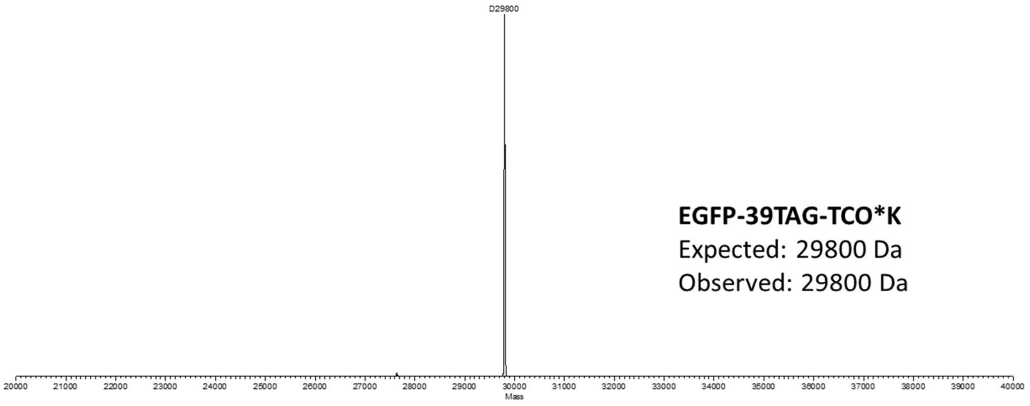
D29800
EGFP-39TAG-TCO*K
Expected: 29800 Da
Observed: 29800 Da
Mass

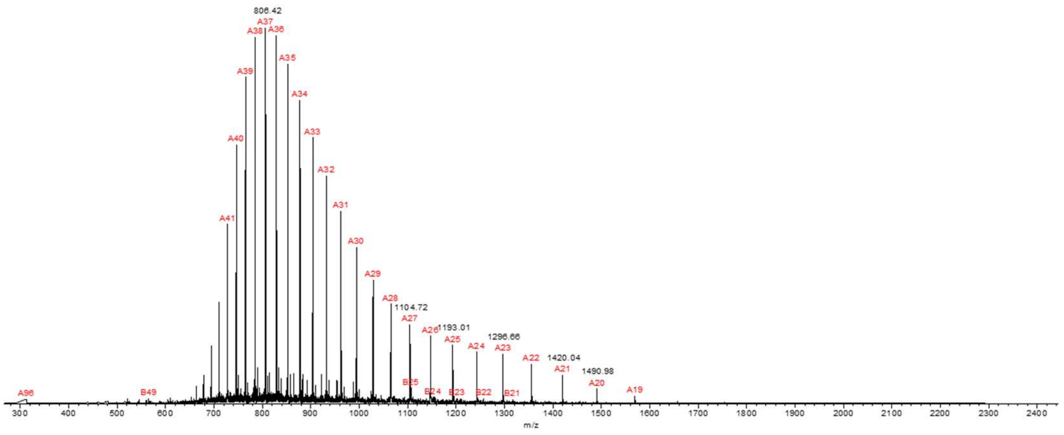
806.42
1104.72
1193.01
1296.66
1420.04
1490.98
m/z

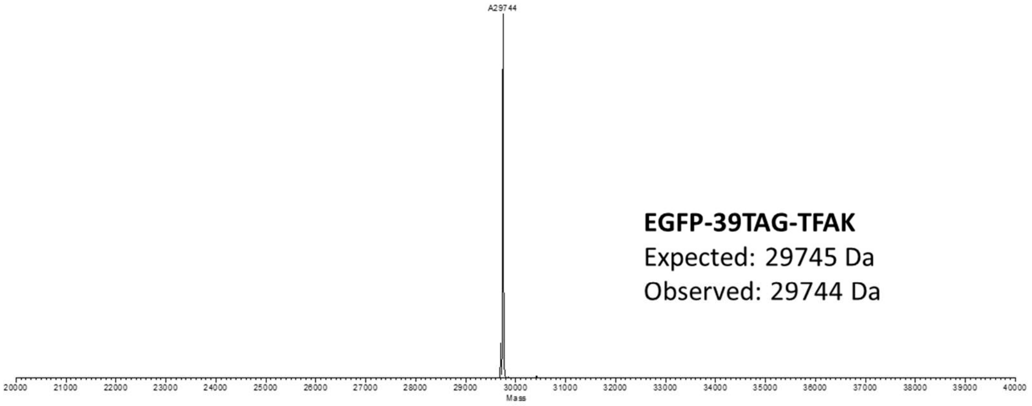

A29744
EGFP-39TAG-TFAK
Expected: 29745 Da
Observed: 29744 Da
Mass


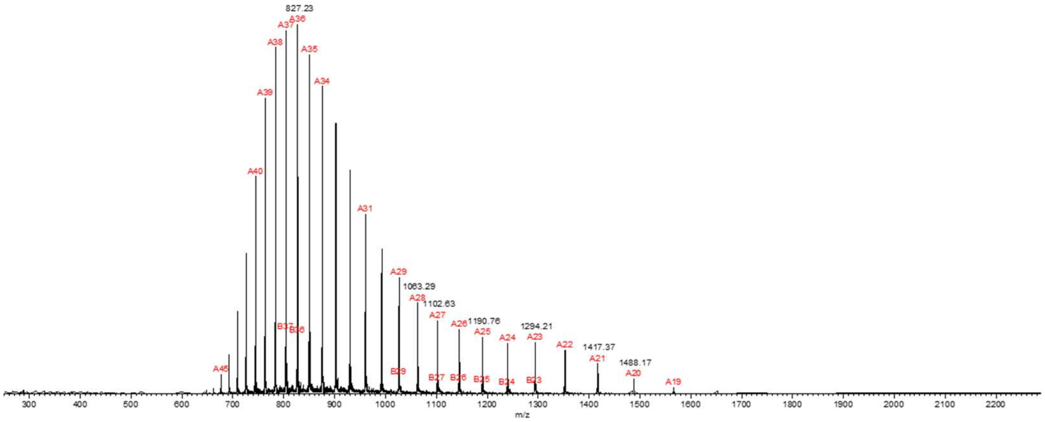

827.23
1063.29
1102.63
1190.76
1294.21
1417.37
1488.17
m/z


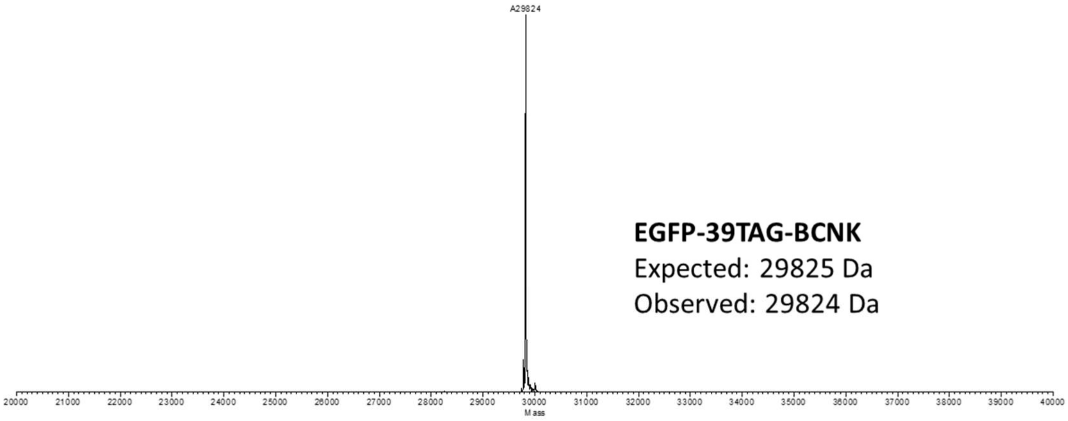

A29824
EGFP-39TAG-BCNK
Expected: 29825 Da
Observed: 29824 Da
Mass


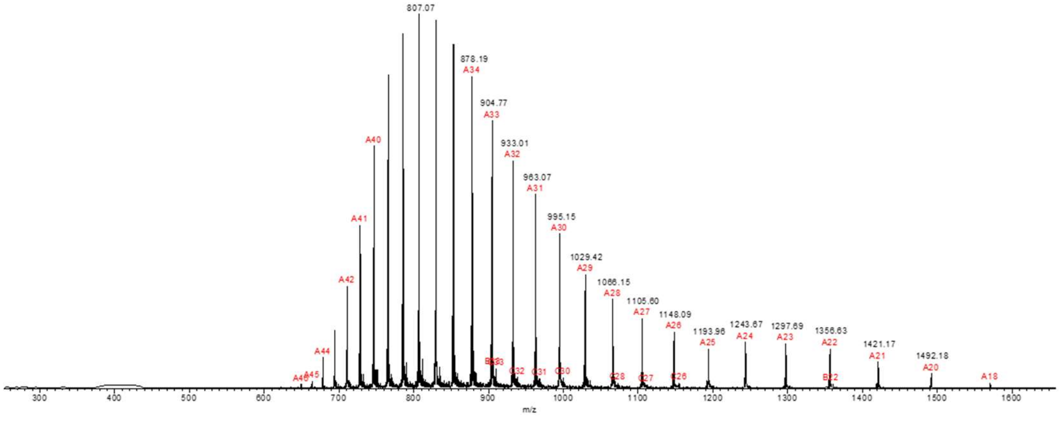

807.07
878.19
904.77
933.01
963.07
995.15
1029.42
1066.15
1105.60
1148.09
1193.96
1243.67
1297.69
1356.63
1421.17
1492.18
m/z

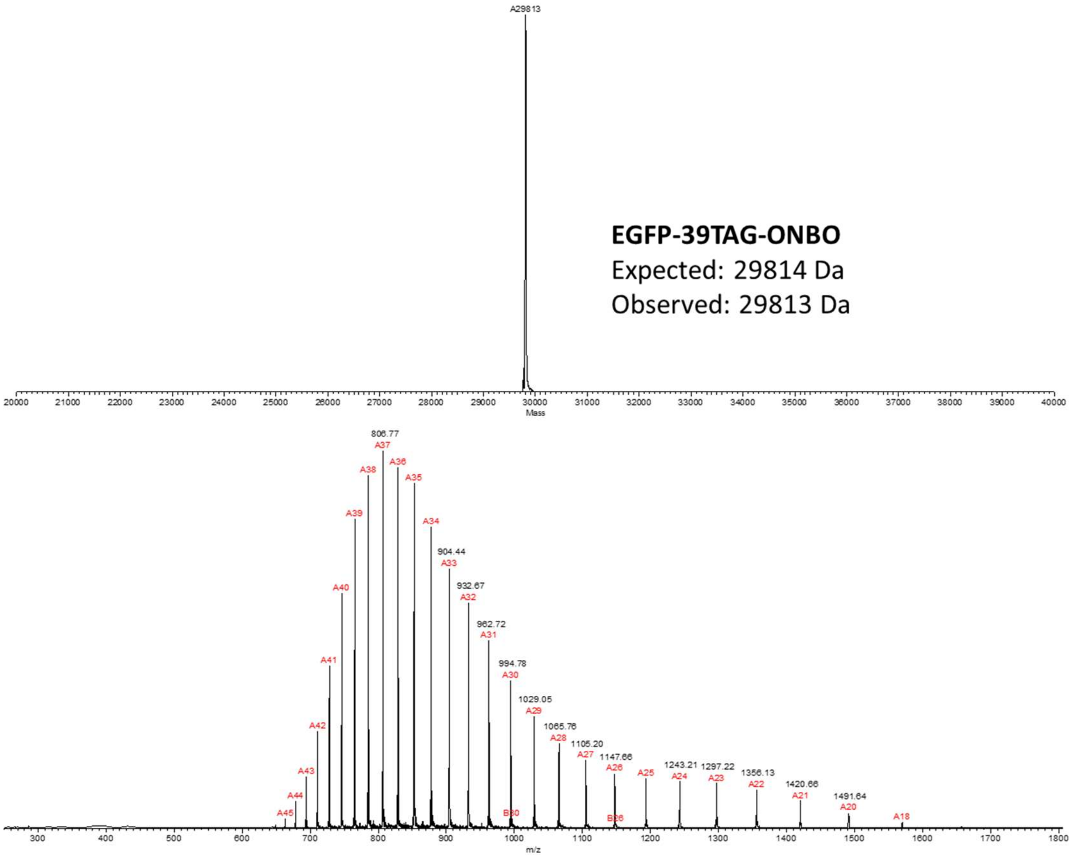
A29813
EGFP-39TAG-ONBO
Expected: 29814 Da
Observed: 29813 Da
Mass
m/z

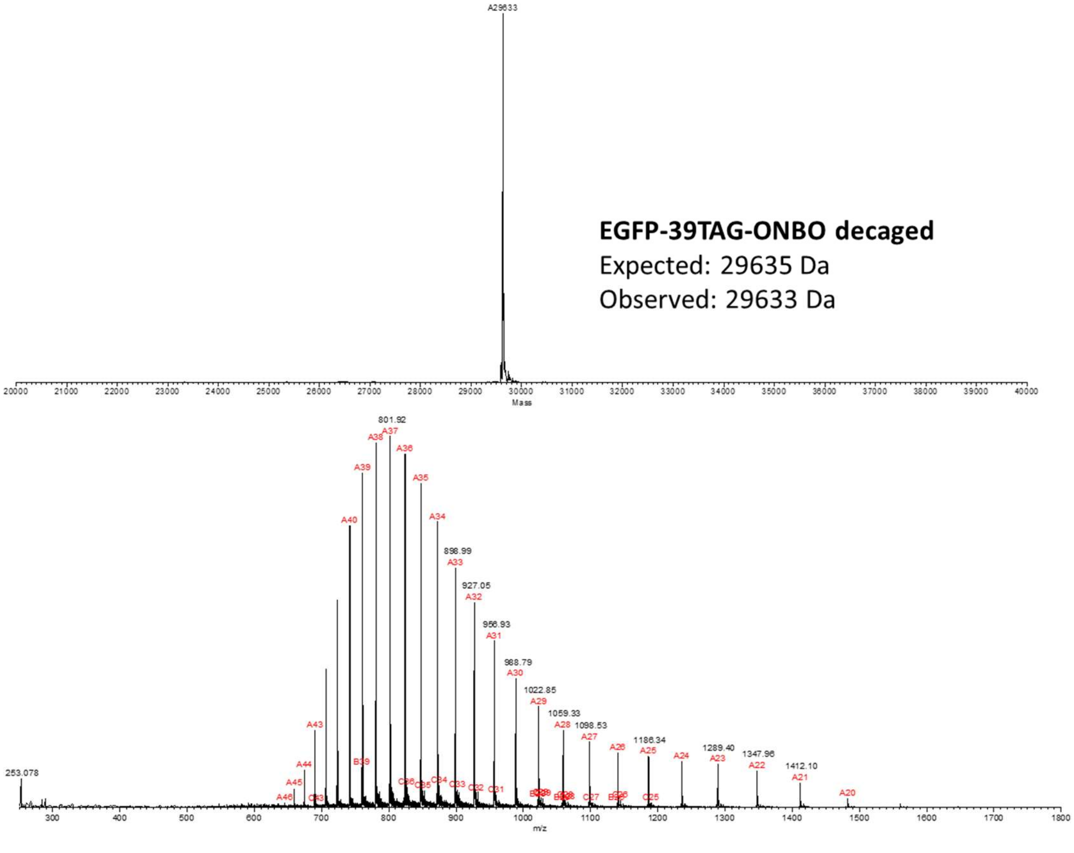
A29633
EGFP-39TAG-ONBO decaged
Expected: 29635 Da
Observed: 29633 Da
Mass
m/z

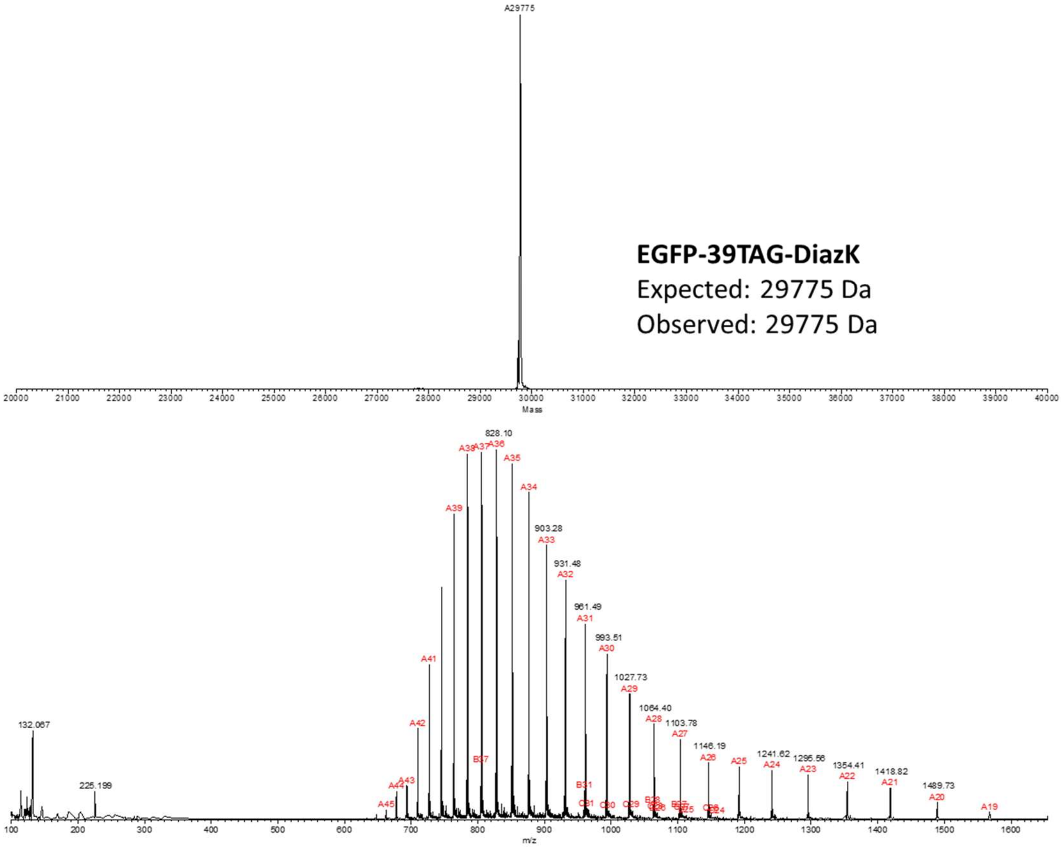


**EGFP-39TAG-DiazK**
Expected: 29775 Da
Observed: 29775 Da

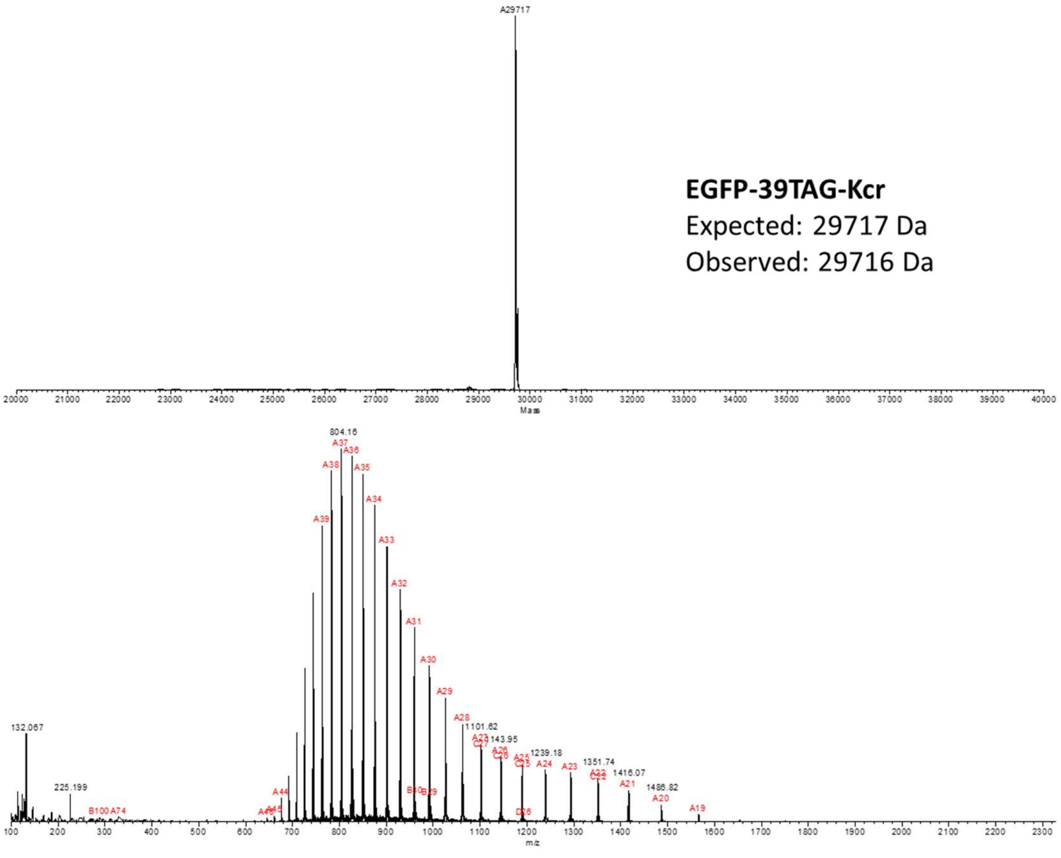


**EGFP-39TAG-Kcr**
Expected: 29717 Da
Observed: 29716 Da

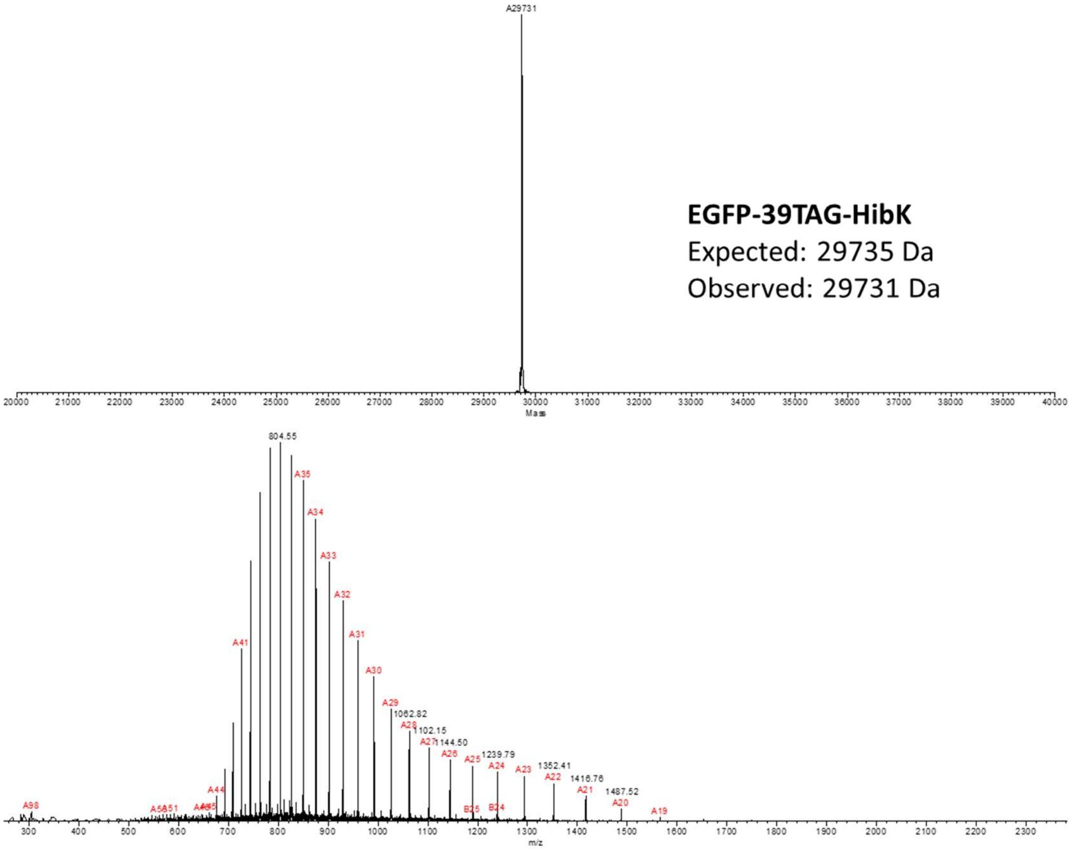
A29731
EGFP-39TAG-HibK
Expected: 29735 Da
Observed: 29731 Da
Mass
804.55
m/z

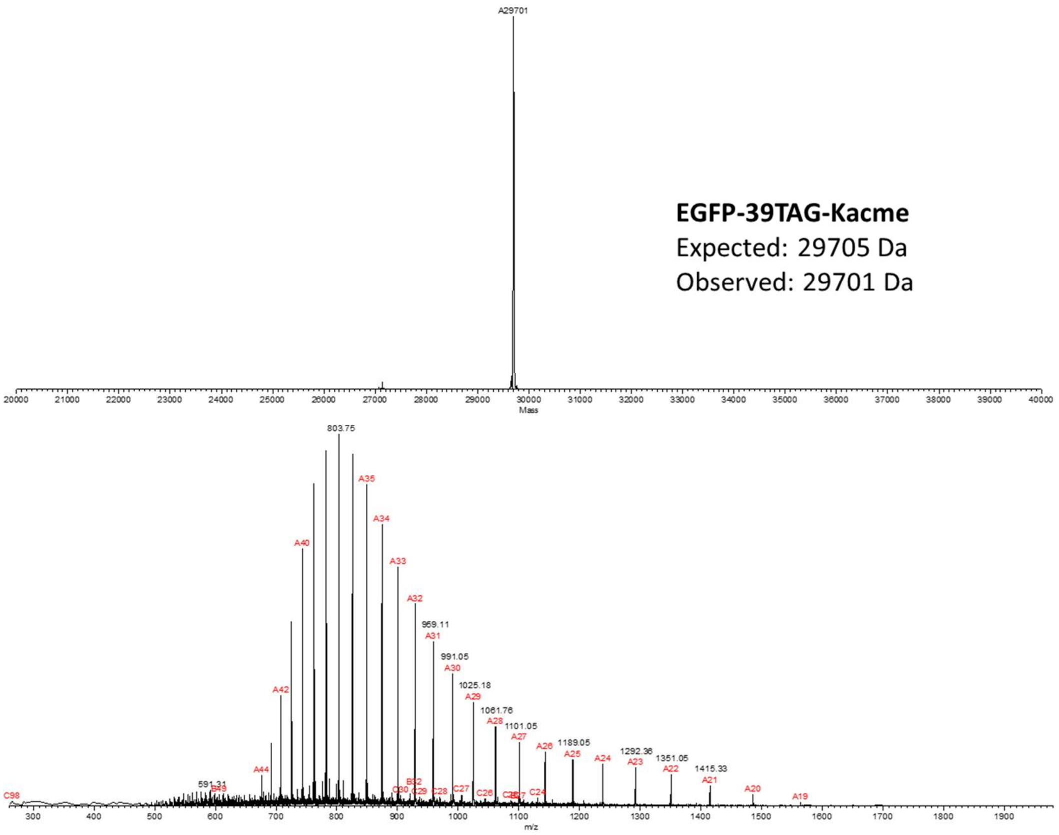
A29701
EGFP-39TAG-Kacme
Expected: 29705 Da
Observed: 29701 Da
Mass
803.75
m/z

**Figure S7**. Deconvoluted and raw ESI-MS spectra of purified EGFP-39-TAG reporters incorporating the designated ncAAs. The expected size of the reporter is noted to the right of each spectrum. Table S5 summarizes the EcLeuRS mutant used in each case and the isolated reporter yield.

**Uncropped gel image from Figure 4G**

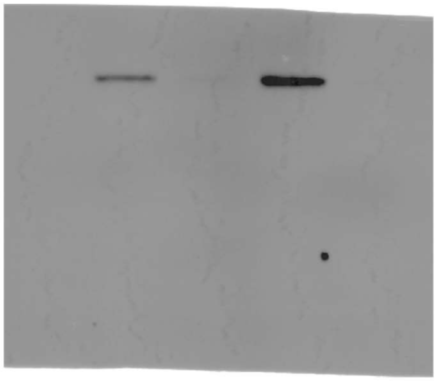

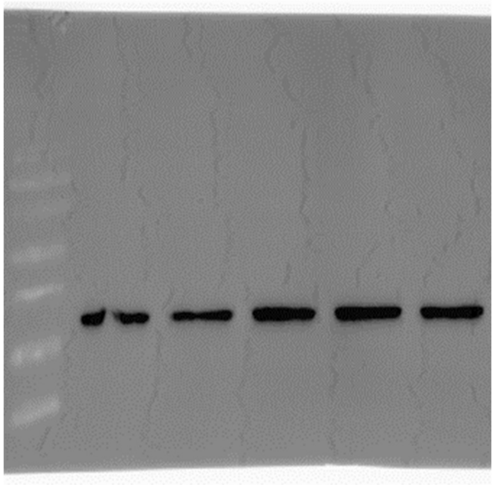

### Methods

#### Materials

All cloning and plasmid propagation was done in DH10B *E. coli* cells. Restriction enzymes were purchased from New England Biolabs (NEB). Polymerase chain reactions (PCR) were carried out using the Phusion High-Fidelity DNA Polymerase from Thermo Fisher. T4 DNA Ligase was purchased from Enzymatics. Plasmid DNA and PCR products were purified using spin columns from Epoch Life Science and Macherey-Nagel Binding Buffer NTI. Sanger sequencing and oligo purchases were completed through GENEWIZ by Azenta Life Sciences. Primer sequences used in this work can be found in Table S1. Antibiotics were purchased from Sigma-Aldrich or Fisher Scientific. Media components were purchased from Fisher Scientific unless otherwise stated. For LB plates and liquid cultures, the following antibiotic concentrations, unless otherwise mentioned were: 100 μg/ml ampicillin.

#### Noncanonical amino acids (ncAAs)

NcAAs were purchased from the indicated companies: Cap **1** (TCI America, Portland, Oregon), AcK **13** (Thermo Scientific Chemicals), AzK **8** (Iris Biotech GMBH, Germany), Kcr **18** (Chem Impex), TfAcK **14** (Millipore Sigma), TCO*K **10**, TCO4K **11**, BCNK **12**, DiazK **6**, SCOK **9**, and CpK **5** (Sirius Fine Chemicals, SiChem GMBH, Germany). C5Az **4**,[1] IsovalK **19**,[2] PropK **15**,[3] ButyrylK **17**,[3] AcrylK **16**,[4] AllylK **7**,[5] and HibK **20**[6] were synthesized as previously described. The synthesis of Kacme **3**, ONBC **2**, and ONBO **21** are described below.

#### General cloning

DNA oligos were purchased from Integrated DNA Technologies (IDT) and GENEWIZ and are listed in in Table S1. For ligation cloning, correct size digestion products were purified by agarose gel purification and a 50 ng vector ligation was performed with at least 3:1 molar ratio of insert to vector.

EcLeuRS hit cloning into pIDT for transfection testing

EcLeuRS hits that were miniprepped from DH10B *E. coli* cells were PCR amplified from the pESC backbone with primers EcLRS-NheI-F and EcLRS-EcoRI-R, digested with EcoRI/NheI, and ligated into a pIDT backbone.[7,8]

pIDT-4xU6-LeuIGI1-EGFP-39TAG

A pIDT backbone was used as the starting point to incorporate 4xU6-LeuIGI1[9] tRNA copies following a previously described cloning method using NheI/AvrII.[8] The EGFP-39TAG reporter was inserted into the pIDT backbone at an SbfI cut site.

EcLeuRS hit cloning into pIDT-4xLeuIGI1-EGFP-39TAG

EcLeuRS hits were PCR amplified from a pIDT backbone with primers EcLRS-NheI-F and EcoRI-EcLeuRS-iR, digested with NheI/XhoI, and ligated into a pIDT-4xU6-LeuIGI1-EGFP-39TAG backbone.

Histone H4 reporter plasmid

The histone H4 sequence was obtained from plasmid mEmerald-H4-23 Addgene (#54117).[10] First, the K5TAG mutant was made using an overlap extension cloning by amplifying piece A with CMV-NdeI-F and H4-K5TAG-iR, and piece B with primers H4-K5TAG-iF and GFP-Cterm-R. Pieces A and B were purified by agarose gel and then PCR amplified for overlap extension using terminal primers CMV-NdeI-F and GFP-Cterm-R, followed by digestion with NheI/HindIII and ligation into mEmerald-H4-24, resulting in mEmerald-H4-K5TAG. The K12TAG mutant was made using mutagenic primers H4-K12TAG-iR and H4-K12TAG-iF, followed by Gibson incubation (HiFi DNA Assembly Master Mix, NEB), resulting in mEmerald-H4-12TAG. Next, a c-terminal 10xHis tag was cloned into both histone H4 mutants using an overlap extension cloning method by amplifying piece A with primers pAcBac1-Histone-H4-NheI-F and Histone-H4-10xHis-TGA-R, and piece B with primers pAcBac1-10xHis-TGA-F and pAcBac1-XbaI-XhoI-R. Pieces A and B were purified by agarose gel and then PCR amplified for overlap extension using terminal primers pAcBac1-Histone-H4-NheI-F and pAcBac1-XbaI-XhoI-F, followed by digestion with NheI/XhoI and ligation into a pAcBac1 vector,[8,11] resulting in pb1-Histone-H4-K5TAG-10xHis and pb1-Histone-H4-K12TAG-10xHis.

**Yeast culture**

The yeast selection host strain MaV203 and MaV203 pGADGAL4-2TAG were a gift from the Schultz lab.[12,13] Yeast chemically competent cells used for small-scale transformations for propagating plasmids were prepared using Zymo Research Frozen-EZ Yeast Transformation II kits. The preparation of liquid and solid media was performed as described previously.[14] All liquid and solid media were supplemented with penicillin-streptomycin (100 IU and 100 μg/mL, respectively, Cytiva Hyclone).

**Yeast spot plating with GAL4-2TAG reporter**

From a single colony, the MaV203 yeast strain containing pGADGAL4-2TAG and pESC-PLRS1-1xSUP4 plasmids were grown overnight in 5 mL SD-Leu-Trp media containing penicillin-streptomycin at 30 °C with shaking.[12,14] The next day, the cultures were supplemented with or without 1 mM Cap (with for positive selection conditions, without for negative selection) into a fresh 10 mL SD-Leu-Trp culture to a starting $OD_{600}$ of 0.1 containing antibiotics. Cultures were grown shaking at 30 °C for 4-6 hours, and then washed with sterile 0.9% NaCl twice using the same volume as the initial culture size (10 mL), then resuspended in a $10^{th}$ the volume of the initial culture volume (1 mL) and allowed to sit at room temperature for 30 minutes. The cultures were then washed two more times with 1 mL 0.9% NaCl before taking the $OD_{600}$ of the final cultures. The cultures were diluted to $OD_{600}$ 0.1, 0.01, and 0.001 in 0.9% NaCl and then spot plated onto 6-well plates containing the selective solid media, with or without 1 mM Cap. The drops were allowed to dry completely next to an open Bunsen burner and then grown upside-down in a 30 °C incubator until growth appeared (24-48 hours).

**Screening for a third TAG site in GAL4**

Cloning a third TAG site in GAL4

Sites for a third TAG site in GAL4 were selected based on structural information (PDB: 3COQ) and previous work.[13] In total, five positions were chosen for testing as an additional TAG site in GAL4: L3, I13, M32, L79, and I121. These sites were mutated using an overlap extension cloning method by PCR amplifying piece A  and piece B using pGADGAL4-2TAG as the template plasmid. Piece A was amplified with the forward primer ADH1-PacI-F and a reverse primer corresponding to each mutant (GAL4-L3TAG-iR, GAL4_I13TAG_R, GAL4_L32TAG_R, GAL4_M79TAG_R, and GAL4_T121TAG_R), and piece B with primer GAL4-SalI-R and an internal forward primer corresponding to each mutation (GAL4-3TAG-iF, GAL4_I13TAG_F, GAL4_L32TAG_F, GAL4_M79TAG_F, and GAL4_T121TAG_F) using pGADGAL4-2TAG as the template (Table S1). Piece A and B were purified by agarose gel and then PCR amplified for overlap extension using terminal primers ADH1-PacI-F and GAL4-SalI-R, followed by digestion with PacI/SalI and ligation into pGADGAL4-2TAG, resulting in five pGADGAL4-3TAG plasmids with each additional third TAG mutation.

Positive and negative selection spot plating of third TAG site

To assess the stringency of mutants containing a third TAG site, spot plating was performed using the original GAL4-2* system and the five new GAL4-3* mutants in the presence or absence of active EcLeuRS. To perform these assays, each pGADGAL4 plasmid was co-transformed into the MaV203 yeast selection host along with either pESC-PLRS1-1xSUP4 or pESC-NoRS-1xSUP4. MaV203 chemically competent cells were prepared for transformation following the Yeastmaker Yeast Transformation System 2 (Clontech). The resulting transformants were tested under positive selection conditions as described under “Yeast spot plating with GAL4-2TAG reporter”. After washes, the cultures were diluted to $OD_{600}$ 0.1, 0.01, and 0.001 in 0.9% NaCl and then spot plated onto 15 cm petri dishes containing the selection conditions, with or without 1 mM Cap. The drops were allowed to dry completely next to an open Bunsen burner and then grown upside-down at 30 °C until growth appeared (24-48 hours).

**Mock selection using GAL4-3* and GAL4-2* selection systems**

To compare the stringency of the GAL-2* and GAL-3* selection systems we co-transformed MaV203 yeast cells with the following plasmid combinations: 1) pGADGAL4-2TAG and pESC-PLRS1-1xSUP4, 2) pGADGAL4-2TAG and pESC-NoRS-1xSUP4, 3) pGADGAL4-3TAG and pESC-PLRS1-1xSUP4, and 4) pGADGAL4-3TAG and pESC-NoRS-1xSUP4. Single colonies from each plate were grown overnight in 5 mL SD-Leu-Trp media containing penicillin-streptomycin at 30 °C with shaking. The next day, these cultures were used to inoculate fresh 10 mL SD-Leu-Trp cultures containing antibiotics to a starting $OD_{600}$ of 0.1, supplemented with or without 1 mM Cap (with for positive selection conditions, without for negative selection). Cultures were grown with shaking at 30 °C for 4-6 hours, and then washed with sterile 0.9% NaCl as described earlier in the “Yeast spot plating with GAL4-2TAG reporter” section. The cultures were diluted to $OD_{600}$ 0.1 for subsequent steps. For both pGADGAL4-2TAG and pGADGAL4-3TAG, cultures encoding active PLRS1 (active) or no EcLeuRS (inactive) were mixed in a 1:100 (active:inactive) ratio, and plated on positive-selection media (-Leu, -Trp, -Ura +ncAA). The plates were allowed to dry completely next to an open Bunsen burner and then grown upside-down at 30 °C until colonies appeared (24-48 hours). 10 colonies from each plate were randomly picked and assessed for the presence of inactive or active EcLeuRS using colony PCR. Roche Diagnostics HotStart ReadyMix was used for colony PCR, following manufacturer's instructions (Kapa Biosystems), using primers LeuRS-4F and pESC-RS-seq-R that yielded an expected band size of 820 bp when EcLeuRS was present, but none when it was absent.

**EcLeuRS MLYYH library design and construction**

Removal of BsaI sites in pESC vector backbone

To prepare the pESC-EcLeuRS-1xSUP4[9] plasmid for Golden Gate assembly cloning, two BsaI recognition sites were first removed from the pESC backbone via cloning. One BsaI recognition site in the ADH1 promoter was mutated from GGTCTCN to AGTCTCN using PrimeSTAR Max DNA Polymerase (Takara Bio) and mutagenic primers ADH1-GBsaIA-F and ADH1-BsaI-R, resulting in plasmid pESC-EcLeuRS-ADH1-GBsaIA-1xSUP4. The other BsaI recognition site in the ampicillin resistance gene ($Amp^R$) was silently mutated from a GGG codon at residue 239 to GGA using an overlap extension cloning method: piece A was PCR amplified from pESC-EcLeuRS-ADH1-GBsaIA-1xSUP4 using primers NsiI-iF and Amp-GGG239GGA-R, piece B was PCR amplified from the same backbone using primers Amp-GGG239-GGA-F and NheI-R. Piece A and B were purified by agarose gel and then PCR amplified for overlap extension using terminal primers NsiI-iF and NheI, followed by digestion with NsiI/NheI and ligation back into pESC-EcLeuRS-ADH1-GBsaIA-1xSUP4, resulting in pESC-EcLeuRS-ADH1-GBsaIA-ΔBsaI-AmpR-1xSUP4.

Mutating the EcLeuRS editing domain

The T252 editing domain of EcLeuRS was mutated to T252A using an overlap extension cloning method by PCR amplifying piece A with primers ADH1-EcoRI-F and LRS-T252A-iR and piece B with primers LRS-T252A-iF and EcLRS-NotI-R from a pESC-EcLeuRS-1xSUP4 backbone. Piece A and B were purified by agarose gel and then PCR amplified for overlap extension using terminal primers ADH1-EcoRI-F and EcLRS-NotI-R, followed by digestion with EcoRI/NotI and ligation into pESC-EcLeuRS-T252A-ADH1-GBsaIA-ΔBsaI-AmpR-1xSUP4, resulting in pESC-EcLeuRS-ADH1-GBsaIA-ΔBsaI-AmpR-1xSUP4.

Golden Gate Assembly Insert site in EcLeuRS

To make the Golden Gate Assembly insertion sequence in EcLeuRS, sticky ends for BsaI insertion were created after residue F490 and before F544 using an overlap extension cloning: piece A was PCR amplified using primers EcLRS-PstI-F and EcLRS-GGA-iR and piece B was amplified using primers EcLRS-499-iF and EcLRS-NotI-R. Piece A and B were purified by agarose gel and then PCR amplified for overlap extension using terminal primers EcLRS-PstI-F and EcLRS-NotI-R, followed by digestion with EcoRI/NotI and ligation into pESC-EcLeuRS-T252A-ADH1-GBsaIA-ΔBsaI-AmpR-1xSUP4, affording the final Golden Gate Assembly vector pESC-EcLeuRS-T252A-GGA-1xSUP4 vector.

EcLeuRS library cloning into Golden Gate Assembly vector

The EcLeuRS library was cloned in two steps. The first round of cloning mutated the M40 and L41 sites, and the second round mutated the Y499, Y527, and H537 sites. First, the M40 and L41 sites were randomized using overlap extension cloning with 32 primers that introduced the appropriate randomized codons (Table S2). The 32 mutagenic primers (Table S3) were mixed in an equimolar ratio prior to PCR amplification of the pESC- EcLeuRS-T252A-GGA vector. Piece A was PCR amplified with ADH1-EcoRI-F and mutagenic EcLRS-M40L41-R primers

(Table S3), and piece B was PCR amplified with primers EcLRS-40-41-iF and EcLRS-NotI-R. Piece A and B were purified by agarose gel and then PCR amplified for overlap extension using terminal primers ADH1-EcoRI-F and EcLRS-NotI-R, followed by digestion with EcoRI/NotI and ligation into pESC-EcLeuRS-T252A-GGA-1xSUP4 vector, ensuring to afford at least >100-fold library coverage upon transforming the 32 EcLeuRS member library into electrocompetent DH10B *E. coli* cells.

To mutate the Y499, Y527, and H537 sites, the previously made vector containing the 32 randomized M40L41 members was PCR amplified with PrimeSTAR Max DNA Polymerase (Takara Bio) with mutagenic primers (Table S4) to introduce the desired mutated sites flanked by the BsaI sticky ends for Golden Gate Assembly insertion. After PCR amplification, the EcLeuRS library members were isolated using agarose gel purification and Golden Gate Assembly was performed with ~1 μg pESC-EcLeuRS-T252A-GGA-1xSUP4 as the receiving vector, using BsaI-HFv2 for scarless introduction of EcLeuRS library members. Golden Gate Assembly reactions were concentrated by ethanol precipitation with yeast tRNA (Invitrogen) and transformed into electrocompetent DH10B *E. coli*. >$1.0 \times 10^6$ transformants were plated (>100-fold library coverage). The colonies were pooled and the library DNA was miniprepped for transformation into the yeast MaV203 pGADGAL4-3TAG selection host.

**EcLeuRS library transformation in yeast**

pGADGAL4-3TAG was transformed into the MaV203 yeast selection host, affording MaV203 pGADGAL4-3TAG. MaV203 pGADGAL4-3TAG chemically competent cells were prepared for EcLeuRS library transformation following the Yeastmaker Yeast Transformation System 2 (Clontech) according to the library-scale protocol using 15 μg of plasmid library per transformation. A total of six transformations were carried out, affording $2.0 \times 10^6$ total transformants (>200-fold library coverage). The colonies were pooled and made into glycerol stocks containing 30% glycerol in PBS, as previously described.[14]

**Selection and screening of EcLeuRS library members**

<u>Original selection workflow</u>

Rounds of selection of EcLeuRS library members was followed as previously described,[12-14] proceeding through five total rounds of selection (+/-/+/-/+).

<u>New selection workflow using homologous recombination</u>

The first round of positive selection was followed the same way as the original workflow, except additional washing steps were performed before plating on selection plates (as described under Yeast spot plating with GAL4-2TAG reporter section). Surviving colonies were harvested from plates and the plasmid DNA was miniprepped using the ChargeSwitch Plasmid Yeast Mini Kit (Invitrogen). Next, EcLeuRS mutants encoded in these plasmids were amplified using primers that also introduced 55 nucleotides of homology to the pESC vector backbone on either side (pESC-HomRecomb-F and pESC-HomRecomb-R). To generate the insert, eight PCR each were set up containing 5 μL of miniprepped library DNA as the template and using primers pESC-HomRecomb-F and pESC-HomRecomb-R (Table S1; 30 cycles). In parallel, the pESC-EcLeuRS-1xSUP4 vector was digested with EcoRI/NotI. Both the PCR-amplified insert and the digested vector were isolated using gel purification, followed by ethanol precipitation as described above. Electrocompetent cells of MaV203 pGADGAL4-3TAG were prepared for homologous recombination of the insert and vector DNA using a previously described method.[15] Briefly, 400 μL of the prepared electrocompetent cells were transformed with 2.5 μg of digested vector and 8.5 μg of insert, recovered in a 1:1 mixture of YPD: 1 M sorbitol, then pelleted (1,000 x g 10 minutes) and resuspended in SD-Leu-Trp to a starting $OD_{600}$ of 0.3. The transformed cells were further allowed to grow overnight with shaking at 30 °C, and then were plated on the negative round of selection (SD-Leu-Trp + 0.1% 5-FOA). The surviving colonies were harvested, the associated plasmid was isolated, and the encoded EcLeuRS was amplified for another round of homologous recombination-mediated introduction into fresh host cells containing pGADGAL4-3TAG. This time, after homologous recombination, the recovered cells were grown overnight in the presence of 1 mM ncAA and then plated on the third and final round of selection (SD-Leu-Trp-Ura +/- 1 mM ncAA) after several washes. 96 of the surviving colonies were screened for ncAA-dependent LacZ expression using a blue/white screen as previously described.[2,12]

**Mammalian cell culture**

HEK293T (ATCC) cell cultures were maintained at 37 °C and at 5% $CO_2$ in DMEM-high glucose (HyClone) supplemented with 10% fetal bovine serum (Corning) and 100 U/mL antibiotic antimycotic penicillin/streptomycin/fungiezone (HyClone).

**EcLeuRS hit testing by transfection**

Hit testing analysis was conducted by transfecting HEK293T cells in 12-well plates with 0.5 μg each of a constructed hit pIDT-EcLeuRS plasmid, pAAV-ITR-CMV-mCherry-1xU6-LeuIGI1, and pAcBac1-CMV-EGFP-39TAG added in the presence and absence of ncAA. Transfections were done with polyethylenimine (PEI, Sigma Aldrich NC0704680) and DNA which was mixed at a ratio of 3.5 μL PEI (1 mg/mL) to 1 μg DNA in DMEM (17.5 μL per 1 μg of DNA), and incubated for 10 min. Fluorescence images and fluorescence protein expression analysis were performed two days post-transfection. Images of each well were taken on Zeiss Axio Observer A1 microscope on brightfield, GFP and mCherry channels. Fluorescence analysis was obtained for both EGFP and mCherry by removal of media and cell lysis in 300 μL with CelLytic M buffer (Sigma) with 0.01 μL/mL Pierce universal nuclease (Fisher). Lysis solution was then incubated for 10 min while nutating at room temperature, then lysate was transferred into a clear bottom 96-well assay plate (Corning, Corning, NY). EGFP fluorescence was recorded at 488 nm excitation and 520 nm emission while mCherry fluorescence was recorded using 588 nm excitation and 620 nm emission using a BioTek Synergy Neo2 Hybrid Multimode Reader. The fluorescence values for untransfected wells were treated as background level fluorescence and were subtracted from each sample. Then EGFP fluorescence for each sample was normalized relative to its mCherry fluorescence (internal control).

**EGFP-39TAG reporter protein expression and purification with EcLeuRS hits**

7.5 million HEK293T cells/plate were seeded in 10 cm cell-culture dishes, and then transfected the next day at 70-80% confluence with 10 µg of pIDT-EcLeuRS-4xLeuIGI1-EGFP-39TAG-6xHisTag. 1 mM of appropriate ncAA was also added to the cells at this time. To increase protein production, plasmids were transfected using 50 µL polyethylenimine (PEI) MAX and 2 mM sodium butyrate was added to the media. Two days after transfection, the media was removed from the plate and cells were scraped off culture dishes with 2 mL 1xPBS. The cells were then spun down at 5,000 x g for 8 min at room temperature, and the supernatant was discarded with the cell pellet stored at -80 °C until purification. For purification, the cell pellet was lysed with 600 µL cell lysis buffer containing celLytic M, protease inhibitor and universal nuclease solution at a 10,000:100:1 ratio, and then nutated at room temperature for 20 minutes. The cell-free extract was then centrifuged for 10 minutes 18,000 x g and the supernatant from the total cell lysate was transferred to new tubes. 2x volume of equilibrium buffer containing 20 mM Na2HPO4, 300 mM NaCl, pH 7.4, 25 mM Imidazole, pH 7.4, was added before loading lysate onto a HisPur Ni-NTA resin (Thermo Scientific) and purified following the manufacturer's protocol. Protein purity was confirmed by SDS-PAGE analysis, and protein molecular mass was characterized using ESI-MS (Agilent Technologies, 1260 Infinity ESI–TOF).

### Histone H4-Kacme expression

HEK293T cells/plate were seeded in 6-well cell culture dishes, and then transfected the next day at 70-80% confluence with 1 µg each of pIDT-EcLeuRS-EF9-4xLeuIGI1 and pAcBac1-histone-H4-K5TAG-10xHis or pAcBac1-histone-H4-K12TAG-10xHis using PEI-MAX. 1 mM of Kacme was added to the cells at the same time for the +ncAA conditions. Two days after transfection, the media was removed from the plate and cells were scraped off culture dishes with 1 mL 1xPBS. The cells were harvested by centrifugation at 5,000 x g for 8 min at room temperature, and the cell pellet stored at -80 °C until lysis.

For lysis, cell pellets were resuspended in 100 µL 1% SDS/ PBS solution and sonicated for two rounds of 10 pulses at 60% amplitude at 4°C, followed by centrifugation at 20,000 g for 10 min. Clarified lysate concentrations were then normalized to 2 mg/mL and the samples were resolved using a 10% SDS-PAGE gel. Proteins were transferred to a PVDF membrane (Cytiva) using a Trans-Blot Turbo transfer system (Bio-Rad). After transfer, membranes were incubated with blocking solution (5% nonfat dry milk in 0.1% Tween 20 TBS buffer) for 3 hours at room temperature. After blocking, membranes were incubated with either Anti-His (Invitrogen, 1:3000) or GAPDH (Invitrogen, 1:1000) primary antibody in fresh blocking solution overnight at 4°C. Following overnight incubation, the primary antibody solution was removed, membranes were washed 5 x 5 min with wash solution (0.1% Tween 20 in TBS), followed by incubation with secondary antibody (chicken anti-mouse HRP, Invitrogen, 1:4000) in blocking solution for 1 hour at room temperature. Membranes were then washed 5 x 5 min, developed using SuperSignal West Dura kit (Fisher), and signal was detected using a ChemiDoc MP imaging system (BioRad).

### Decaging ONBO

0.5 μM purified EGFP-39TAG-ONBO was diluted in 50 mM HEPES containing 5 mM L-ascorbic acid and was irradiated for 10 minutes on ice and in a 4 °C room at 365 nm . After irradiation, the sample was directly analyzed by intact protein ESI-MS.

**Table S1.** Oligonucleotides used in this work

| Oligo name | Sequence (5' to 3') |
|---|---|
| EcLRS-NheI-F | attattagctagcgccaccatggaagagcaataccgcccg |
| EcLRS-EcoRI-R | attattagaattcttaaacgggcccgccaacgac |
| EcoRI-EcLeuRS-iR | agactcgagttaaagtcgacttaacgcgttgaattcttaaacgggcccgccaacg |
| ADH1-GBsaIA-F | taataccttcgttagtctccctaacatgtaggtggcggag |
| ADH1-GBsaIA-R | tgttagggagactaacgaaggtattataggaatcccgatgtatggg |
| NsiI-iF | attattaatgctttgttagaaaaatagcgctctcggg |
| Amp-GGG239GGA-R | taccgcgagatccacgctcaccggctccagatttatc |
| Amp-GGG239GGA-F | ggtgagcgtggatctcgcggtatcattgcagcactg |
| NheI-R | attattataccatcatttcttcgatcccgaccgg |
| ADH1-EcoRI-F | attattatcaagctataccaagcatacaatcaactg |
| LRS-T252A-iR | gtacaacccataaacgcgtccgggcgggtagtgtaaacg |
| LRS-T252A-iF | cgcccggacgcgtttatgggttgtacctacctggcgg |
| EcLRS-NotI-R | aataatgcggccgcttaaacgggcccgccaacgaccag |
| EcLRS-PstI-F | attattacgtatgctgggcaaaaacgtc |
| EcLRS-GGA-iR | tcacgcatcagtttgtggaagaagtgagacctacgtaaggtctccggtgtcgaaagtgtcggt ttcac |
| EcLRS-GGA-iF | cttcttccacaaactgatgcgtga |
| EcLRS-40-41-iF | ccctatccttctggtcgactacac |
| pESC-HomRecomb-F | tttctttttctgcacaatatttcaagctataccaagcatacaatcaactgaattcatggaagagca ataccgcccg |
| pESC-HomRecomb-R | agagctcagatcttatcgtcgtcatccttgtaatccatcgatactagtgcggccgcttaaacgg gcccgccaac |
| ADH1-PacI-F | attattagaagggaactttacacttctcctatgcac |
| GAL4-SalI-R | attattaatgatatatggtgggacctgttgtggtac |

| | |
|---|---|
| GAL4-3TAG-iF | ctgaaagatgaagtagctgtcttctatcgaacaagcatgcg |
| GAL4-L3TAG-iR | cgatagaagacagctacttcatctttcaggaggcttgcttc |
| GAL4_I13TAG_F | gcatgcgattagtgccgacttaaaaagctcaagtgc |
| GAL4_I13TAG_R | aagtcggcactaatcgcatgcttgttcgatagaagac |
| GAL4_L32TAG_F | gcgccaagtgttagaagaacaactgggagtgtcgctac |
| GAL4_L32TAG_R | cagttgttcttctaacacttggcgcacttcgg |
| GAL4_M79TAG_F | gagaagaccttgactagattttgaaaatggattctttacaggatataaaagc |
| GAL4_M79TAG_R | gaatccattttcaaaatctagtcaaggtcttctcgaggaaaaatcagtag |
| GAL4_T121TAG_F | ctgatatgcctctatagttgagacagcatagaataagtgcgac |
| GAL4_T121TAG_R | ctatgctgtctcaactatagaggcatatcagtctccactgaag |
| pESC-RS-seq-R | ctcgttccctttcttccttgtttc |
| LeuRS-4F | ctcgttccctttcttccttgtttc |
| 2CYC-seq-R | gaagaatgagccaagacttgc |
| CYC-term-F | agagcgcccaatacgcaaac |
| CMV-NdeI-F | caagtgtatcatatgccaagtacgccccctattg |
| H4-K5TAG-iR | cttcccgccctagccgcggccagacatggtgg |
| H4-K5TAG-iF | gccgcggctagggcgggaagggtcttggcaaag |
| GFP-Cterm-R | cggtgaacagctcctcgcccttgctcac |
| H4-K12TAG-iR | gcgccgccctagccaagacccttcccgcctttg |
| H4-K12TAG-iF | ggtcttggctagggcggcgctaagcgccaccg |
| pAcBac1-Histone-H4-NheI-F | attattagctagcgccgccaccatgtctggccgcgg |
| Histone-H4-10xHis-TGA-R | gaattctcaatggtgatgatgatggtggtgatgatgatgcccaccgaaaccgtagaggg |

| | |
|---|---|
| pAcBac1-10xHis-TGA-F | gcatcatcatcaccaccatcatcatcaccattgagaattcaacgcgttaagtcgacaatc |
| pAcBac1-XbaI-XhoI-R | acgggccctctagactcgag |

**Table S2.** Library design of residues randomized in EcLeuRS

| EcLeuRS site | Mutants (codon) |
|---|---|
| M40 | G (GGC), A (GCA), V (GTT), I (ATT) |
| L41 | S (TCA), P (CCC), T (ACG), E (GAG), Q (CAA), N (AAC), L (CTT), V (GTA) |
| T252 | A (GCG) |
| Y499 | G (GGC), S (TCA), R (CGT), A (GCT), I (ATC), L (TTG), V (GTA) |
| Y527 | NBK |
| H537 | G (GGA), T (ACT) |

N=A/C/G/T; B=C/G/T; K=G/T

**Table S3.** M40L41 mutagenic primers for EcLeuRS library cloning

| Oligo name | Sequence (5' to 3') |
|---|---|
| EcLRS-M40L41-R | gtgtagtcgaccagaaggatagggNNNnnnagacaggcagtaatacttctctttgc |

NNN=GGC/GCA/GTT/ATT; nnn=TCA/CCC/ACG/GAG/CAA/AAC/CTT/GTA

**Table S4.** Y499, Y527, and H537 mutagenic primers for EcLeuRS library cloning

| Oligo name | Sequence (5' to 3') |
|---|---|
| EcLRS-BsaI-Y499-F | attattaggtctcc**cacc**tttatggagtcctcctgg**NNNt**atgcgcgctacacttgccc |
| EcLRS-BsaI-Y527NBK-H537G-R | taataatggtctcc**gaag**cggaagtagagcagccccataatggcgtgttcaataccaccaat**mvn**gatatccaccggcagccagtag |
| EcLRS-BsaI-Y527NBK-H537T-R | taataatggtctcc**gaag**cggaagtagagcagcgtcataatggcgtgttcaataccaccaat**mvn**gatatccaccggcagccagtag |

NNN= GGC/TCA/CGT/GCT/ATC/TTG/GTA; mvn= NBK; cacc=BsaI sticky end; gaag=BsaI sticky end

**Table S5. Yield and MS analysis for reporter proteins.** Related to Figure 4. The EGFP-39TAG reporter was expressed in HEK293T cells in the presence of EcLeuRS hits with an engineered $tRNA_{CUA}^{EcLeu}$ pair and their substrates.

| Reporter | EcLeuRS hit | ncAA | Yield (μg/ 10 cm dish) | Expected mass (Da) | Observed mass (Da) |
|---|---|---|---|---|---|
| EGFP-39TAG | EF1 | ONBC | 18 | 29813 | 29812 |
| | B11 | ONBO | 16 | 29814 | 29813 |
| | B11 | ONBO after decaging | - | 29635 | 29633 |
| | EF9 | Kacme | 16 | 29705 | 29701 |
| | B11 | BCNK | 20 | 29825 | 29824 |
| | B11 | DiazK | 18 | 29775 | 29775 |
| | EF9 | TfAcK | 27 | 29745 | 29744 |
| | EF9 | SCOK | 17 | 29798 | 29798 |
| | EF9 | TCO*K | 6 | 29800 | 29800 |
| | B11 | Kcr | 19 | 29717 | 29716 |
| | EF9 | Khib | 12 | 29735 | 29731 |
| EGFP-WT | - | - | 58 | 29684 | 29683 |

**Synthesis of ncAAs**

Synthesis of photocaged citrulline (ONBC **2**)

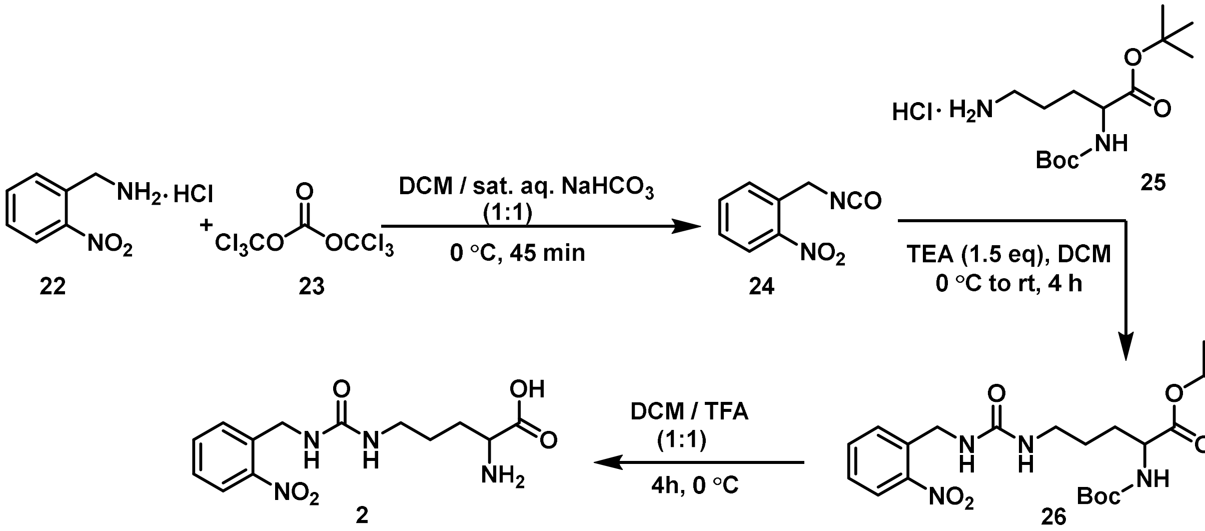


2-nitrobenzylamine hydrochloride salt **22** (1.0 g, 5.3 mmol) was dissolved in a biphasic mixture of $CH_2Cl_2$ (20.0 mL) and sat. aq. $NaHCO_3$ (20.0 mL) and cooled to at 0 ˚C with ice bath. After stirring vigorously for 30 min, a solution of triphosgene **23** (1.57 g, 5.3 mmol) in dry $CH_2Cl_2$ (5.0 ml) was added dropwise. This mixture was vigorously stirred at 0 °C for 15 min then the organic layer was separated and the aq. layer was extracted with 2x10 mL $CH_2Cl_2$. The combined organic layer was washed with brine (10.0 mL), dried over anhydrous $Na_2SO_4$ and concentrated under reduced pressure using a rotary evaporator to give the isocyanate intermediate **3** as an oil (820 mg, 87% yield) which was used directly in the next step without further purification.

To a solution of *N*-Boc-*tert*-butylester of L-ornithine hydrochloride **25** (415 mg, 1.28 mmol, 1.0 equiv.) in $CH_2Cl_2$ (15 mL), $Et_3N$ (270 µL) was added dropwise and cooled to 0 ˚C while stirring. A solution of the intermediate isocyanate **24** (228 mg, 1.0 equiv.) in $CH_2Cl_2$ (5.0 mL) was added and the resulting mixture was stirred at room temperature for 4 h. The organic layer was washed with 2×10 mL water, followed by brine (5.0 mL) and dried over anhydrous $Na_2SO_4$. After concentrating the organic layer under reduced pressure using a rotary evaporator, the resulting crude product was purified by flash chromatography (1:1 EtOAc-hexane; 230-400 silica gel) to afford the *N*-Boc-*tert*-butyl ester of citrulline **26** as an oil (405 mg, 68% yield). **$^1$H NMR (500 MHz, CDCl$_3$):** δ 8.00-7.98 (m, 1H; Ar-H), 7.66-7.64 (m, 1H; Ar-H), 7.58-7.55 (m, 1H; Ar-H), 7.41-7.37 (m, 1H; Ar-H), 5.55 (s,1H; N-H), 5.15-5.14 (m, 2H; 2 N-H), 4.59 (d, *J* = 6.2 Hz, 2H; Ar-CH$_2$), 4.13-4.08 (m, 1H; C-H), 3.19-3.16 (m, 2H; N-CH$_2$), 1.78-1.49 (m, 4H; CH$_2$-CH$_2$), 1.43 (s, 9H; *tert*-Butyl), 1.40 (s, 9H; *tert*-Butyl); **$^{13}$C NMR (125 MHz, CDCl$_3$):** δ 171.8 (C=O), 158.3 (C=O), 155.8 (C=O), 148.3 (C-

$NO_2$), 135.5 (C), 134.0 (CH), 131.9 (CH), 128.3 (CH), 125.0 (CH), 82.2 (C), 80.0 (C), 53.7 (CH), 42.0 ($CH_2$), 40.0 ($CH_2$), 30.7 ($CH_2$), 28.4 ($CH_3$), 28.1 ($CH_3$). HRMS (ESI-TOF) m/z: $[M + H]^+$ Calcd for $C_{22}H_{35}N_4O_7$ 467.2506; found: 467.2617.

The intermediate **2** (250 mg) was dissolved in $CH_2Cl_2$ (4.0 mL) and anhydrous trifluoroacetic acid (4.0 mL) was added dropwise at 0 °C and the resulting mixture was stirred at 0 °C for 4 h to have complete removal of Boc protection (monitored by LC-MS). The mixture was concentrated under reduced pressure. To the resulting oil, DCM (~10 mL) was added and concentrated to dryness. The process was repeated several times to get rid of dissolved trifluoroacetic acid. The expected product **2** (photocaged citrulline)[15] was obtained as a white solid (166 mg) in quantitative yield. **$^1$H NMR (500 MHz, $D_2O$):** δ 8.09 (dd, *J* = 8.2, 1.3 Hz, 1H; Ar-H), 7.74-7.71 (m, 1H; Ar-H), 7.58 (dd, *J* = 7.9, 1.3 Hz, 1H; Ar-H), 7.54-7.50 (m, 1H; Ar-H), 4.62 (s, 2H; Ar-$CH_2$), 3.24-3.21 (m, 1H; C-H), 3.14-3.11 (m, 2H; N-$CH_2$), 1.64-1.45 (m, 4H; $CH_2$-$CH_2$); **$^{13}$C NMR (125 MHz, $D_2O$):** δ 183.4 ($CO_2H$), 160.4 (C=O), 147.5 (C-$NO_2$), 134.9 (C), 134.3 (CH), 129.2 (CH), 128.2 (CH), 125.1 (CH), 55.7 (CH), 41.2 ($CH_2$), 39.7 ($CH_2$), 31.9 ($CH_2$), 25.7 ($CH_2$). HRMS (ESI-TOF) m/z: $[M + H]^+$ Calcd for $C_{13}H_{19}N_4O_5$ 311.1355; found: 311.1380.

<u>Synthesis of photocaged ornithine (ONBO **21**)</u>

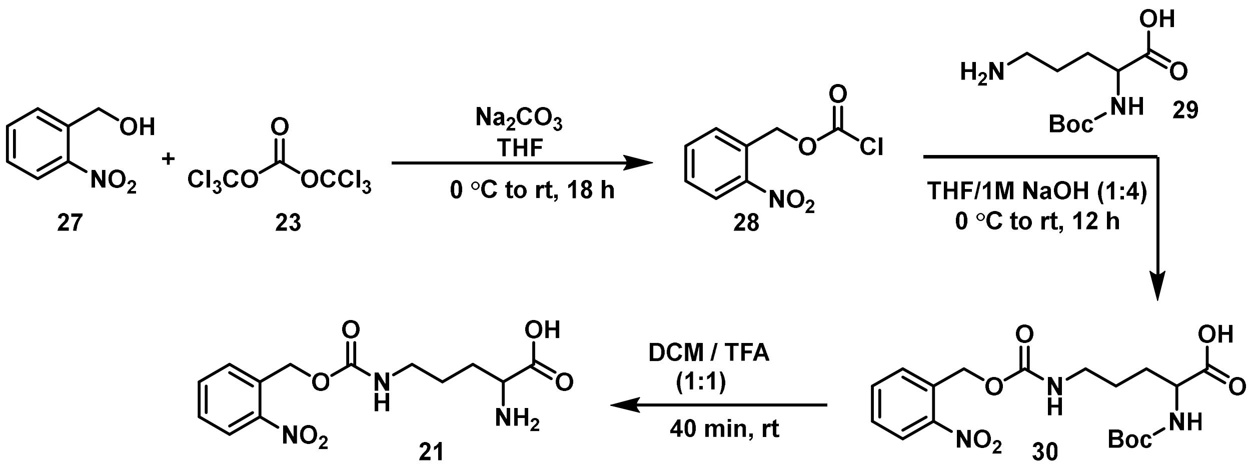


2-Nitrobenzyl alcohol **27** (400 mg, 2.61 mmol) was dissolved in 5.0 mL of dry THF, followed by the addition of $Na_2CO_3$ (277 mg, 2.61 mmol, 1.0 equiv.). After stirring for 5 minutes, the mixture was cooled to 0 °C using an ice bath. A solution of triphosgene **23** (775 mg, 2.61 mmol, 1.0 equiv.) in 3.0 mL of dry THF was then added dropwise. The resulting mixture was stirred overnight while slowly raising the temperature to room temperature. The solvent was removed under reduced pressure using a rotary evaporator (without applying any heat), affording the 2-nitrobenzyl carbonochloridate **28** as a white solid in quantitative yield, which was utilized directly in the subsequent step without further purification.

To a solution of N-Boc-L-ornithine **29** (606 mg, 2.61 mmol) in 2.0 mL of THF and 8.0 mL of 1M NaOH, at 0°C, the crude 2-nitrobenzyl carbonochloridate **8** (562 mg, 2.61 mmol, 1.0 equiv) was added and stirred for 12 hours while slowly increasing the temperature to room temperature. After completion of the reaction, the aqueous layer was washed with 2×10 mL of diethyl ether ($Et_2O$). Subsequently, the aqueous layer was acidified with 20.0 mL of 1M HCl and extracted with 3×10 mL of ethyl acetate (EtOAc), then dried over anhydrous $Na_2SO_4$. The solvent was removed under reduced pressure using a rotary evaporator, yielding the Boc-protected ornithine intermediate **30** as a white solid (762 mg, 71% yield), which was utilized directly in the subsequent step without further purification.

**$^1$H NMR (500 MHz, $CDCl_3$):** δ 8.07 (d, *J* = 8.2 Hz, 1H; Ar-H), 7.66-7.55 (m, 2H; Ar-H), 7.50-7.43 (m, 1H; Ar-H), 5.49 (s, 2H; Ar-$CH_2$), 5.29-5.25 (m, 1H; N-H), 4.35-4.30 (m, 1H; CH), 3.26-3.20 (m, 4H; $CH_2$), 1.90-1.60 (m, 4H; $CH_2$-$CH_2$); **$^{13}$C NMR (125 MHz, $CDCl_3$):** δ 176.0 ($CO_2H$), 156.3 (C=O), 155.9 (C=O), 147.5 (C-$NO_2$), 133.9 (C), 133.3 (CH), 128.9 (CH), 128.7 (CH), 125.1 (CH), 80.5 (C), 63.5 (C $H_2$), 53.1 (CH), 40.7 ($CH_2$), 29.8 ($CH_2$), 28.4 ($CH_3$), 25.9 ($CH_2$).

The intermediate **30** (350 mg) was dissolved in dry $CH_2Cl_2$ (5.0 mL) and anhydrous trifluoroacetic acid (5.0 mL) was added dropwise at 0 °C and the resulting mixture was stirred at 0 °C for 40 min to have complete removal of Boc protection (monitored by LC-MS). The mixture was concentrated under reduced pressure. To the resulting oil DCM (~10 mL) was added and concentrated to dryness. The process was repeated several times to get rid of dissolved trifluoroacetic acid. The crude product was dissolved in minimum amount of MeOH and precipitated using cold $Et_2O$ to get the expected product **21** (photocaged ornithine) as a white solid (265 mg) in quantitative yield.

**$^1$H NMR (500 MHz, $D_2O$):** δ 8.14 (d, *J* = 8.0 Hz, 1H; Ar-H), 7.76 (t, *J* = 7.6 Hz, 1H; Ar-H), 7.63 (d, *J* = 7.8, 1H; Ar-H), 7.57 (t, *J* = 7.9 Hz, 1H; Ar-H), 5.44 (s, 2H; Ar-$CH_2$), 3.24-3.13 (m, 3H; C-H and $CH_2$), 1.64-1.48 (m, 4H; $CH_2$-$CH_2$); **$^{13}$C NMR (125 MHz, $D_2O$):** δ 183.3 ($CO_2H$), 158.0 (C=O), 146.9 (C-$NO_2$), 134.5 (C), 132.6 (CH), 129.0 (CH), 128.6 (CH), 125.0 (CH), 63.6 ($CH_2$), 55.7 (CH), 40.4 ($CH_2$), 31.9 ($CH_2$), 25.3 ($CH_2$). HRMS (ESI-TOF) m/z: $[M + H]^+$ Calcd for $C_{13}H_{18}N_3O_6$ 312.1196; found: 312.1203.

Synthesis of Acetyl-methyllysine (Kacme):

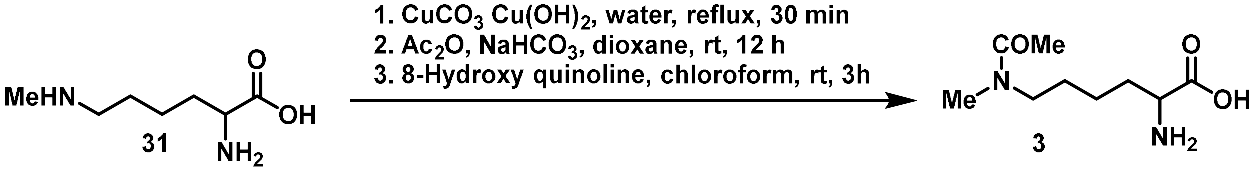


Acetyl-methyllysine **13** was prepared by following the literature procedure[10] and purified by reverse-phase HPLC using a pre-packed C18 column and a water/acetonitrile (supplemented with 0.05% trifluoroacetic acid) gradient as eluent.

**$^1$H NMR (500 MHz, $D_2O$):** (Rotameric mixture) δ 4.05 (q, *J* = 6.5 Hz, 1H; C-H), 3.39-3.32 (m, 2H; N-$CH_2$), 3.01 and 3.32 (2 s, 3H; $CH_3$, rotamer), 2.09 and 2.07 (2 s, 3H; $CH_3$, rotamer), 2.01-1.88 (m, 2H; $CH_2$), 1.49-1.29 (m, 2H; $CH_2$); **$^{13}$C NMR (125 MHz, $D_2O$):** (Rotameric mixtures) major rotamer: δ 174.0 (C=O), 172.0 ($CO_2H$), 60.4 (CH), 50.6 ($CH_2$), 36.1 ($CH_3$), 29.4 ($CH_2$), 26.6 ($CH_2$), 21.3 ($CH_2$), 20.6 ($CH_3$); Minor rotamer: δ 173.8 (C=O), 171.9 ($CO_2H$), 52.6 ($CH_2$), 47.1 ($CH_2$), 33.4 ($CH_3$), 29.3 ($CH_2$), 25.6 ($CH_2$), 20.0 ($CH_3$); HRMS (ESI-TOF) m/z: $[M + H]^+$ Calcd for $C_9H_{19}N_2O_3$ 203.1396; found: 203.1444.

### $^1$H and $^{13}$C NMR Spectra of the Synthesized Compounds:

NMR Characterization of **26**:

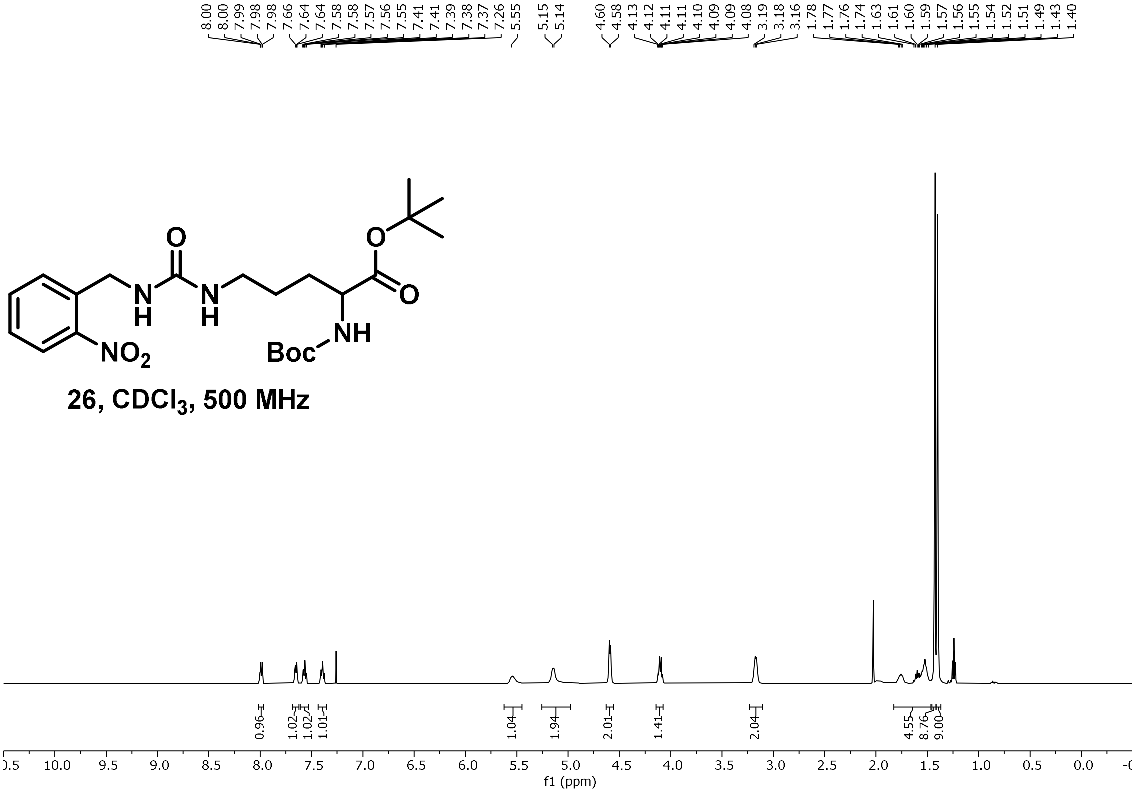


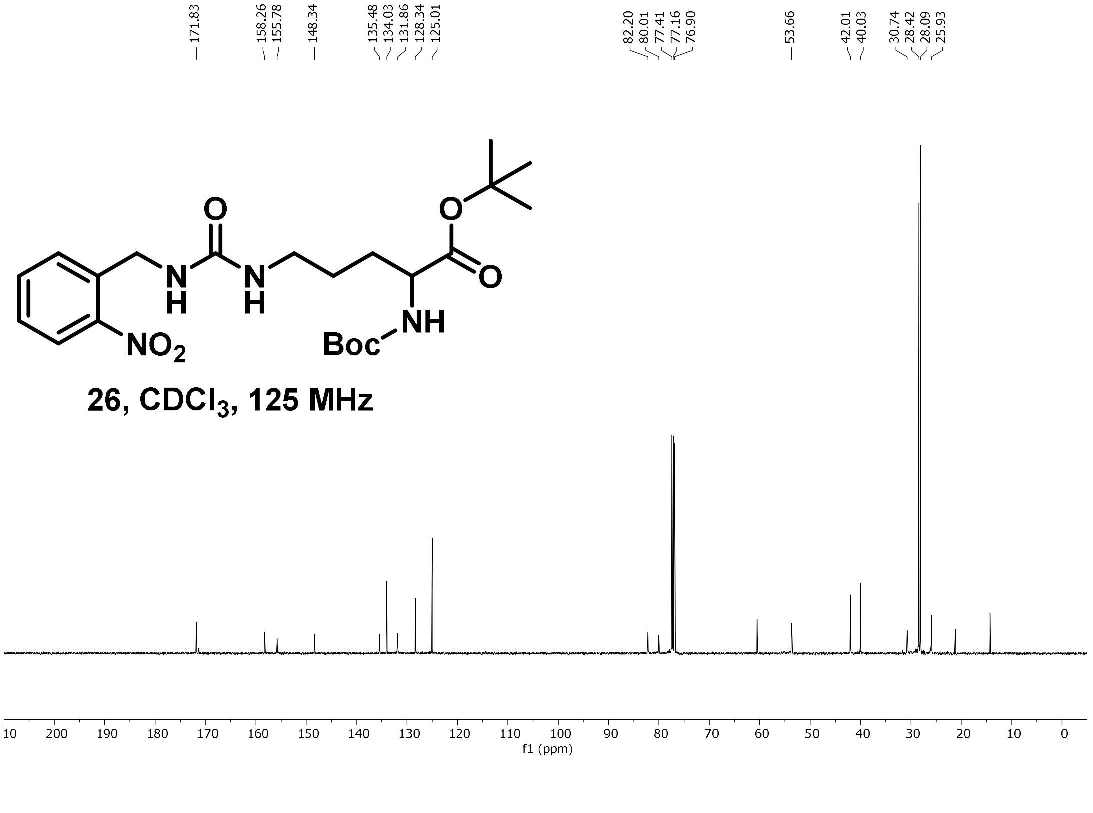

NMR Characterization of (ONBC) **2**:

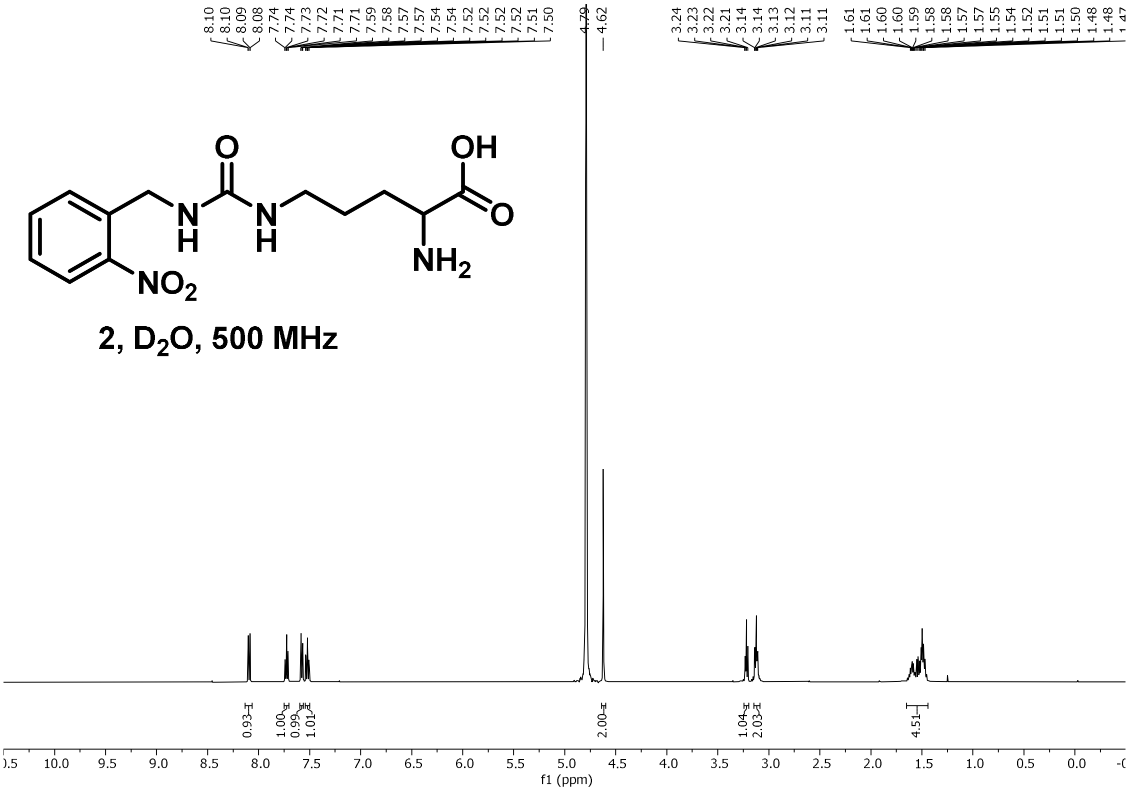


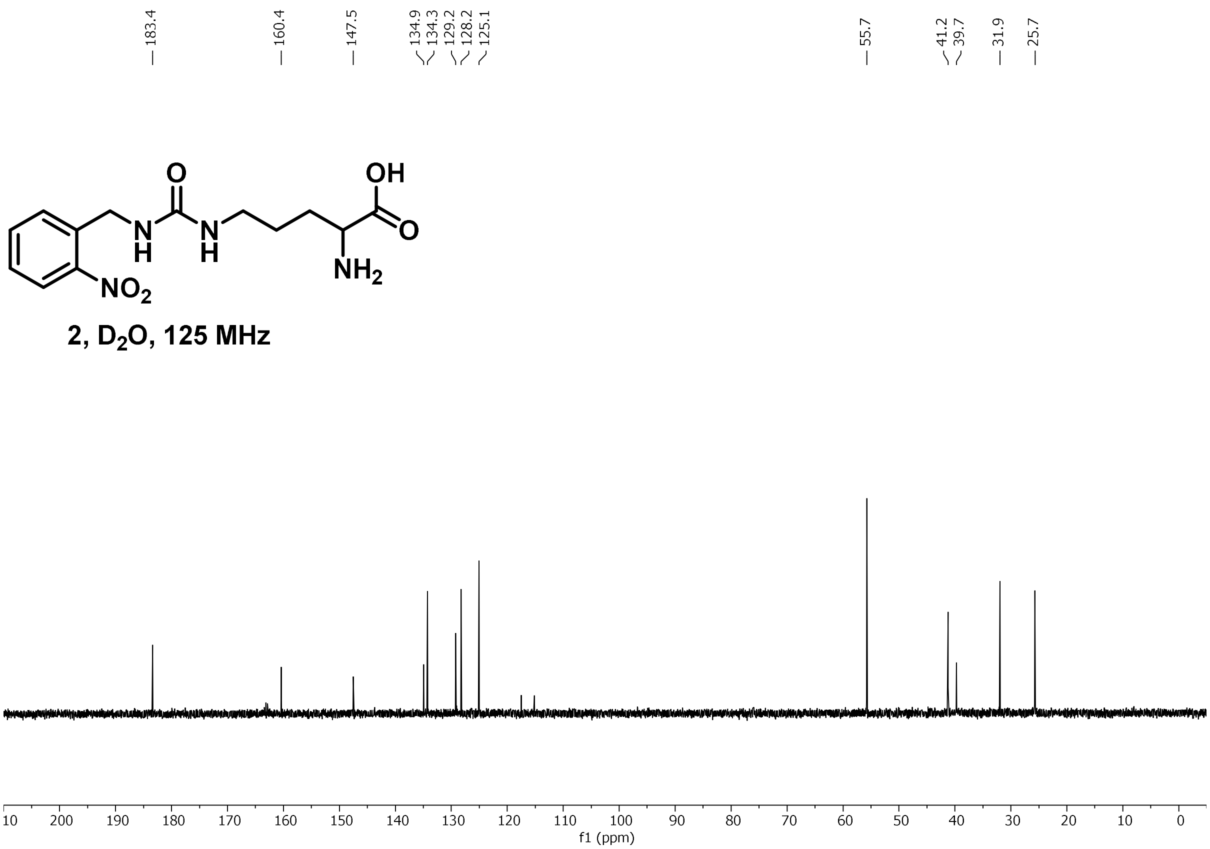

NMR Characterization of **30**:

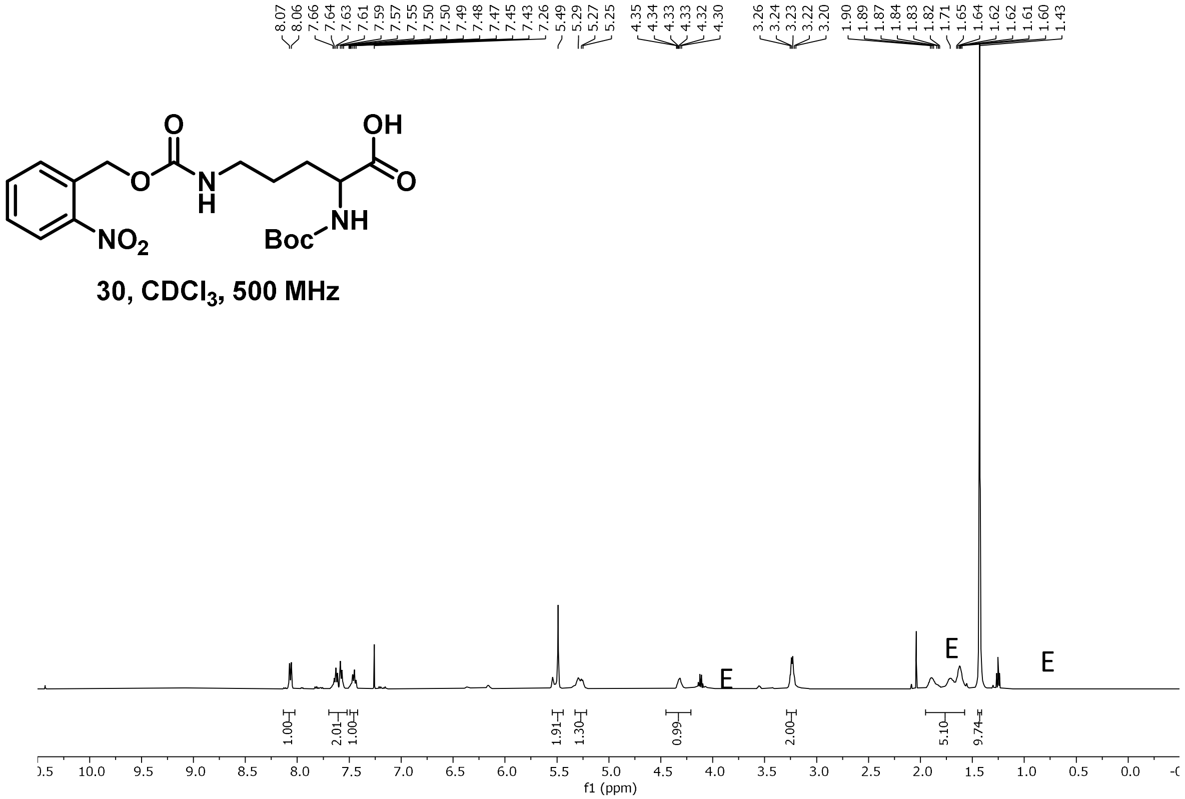


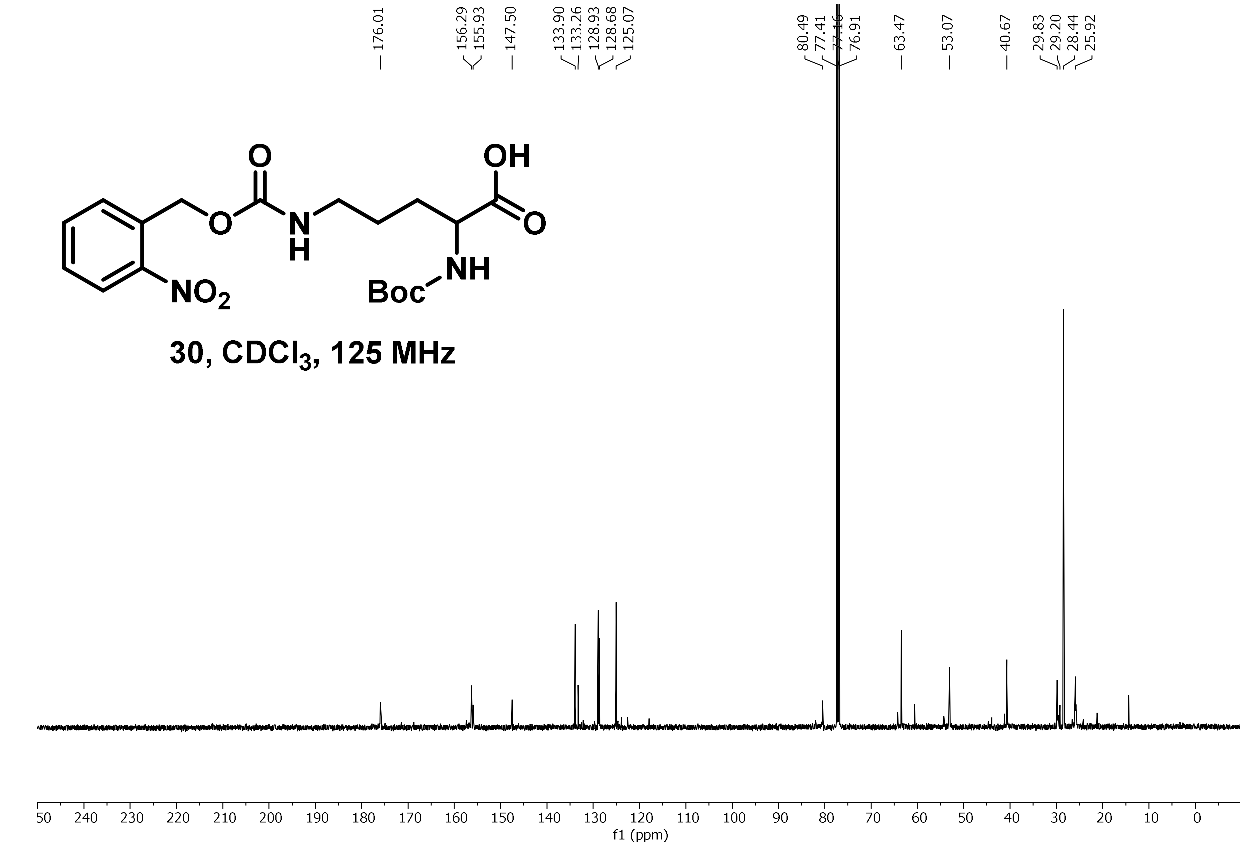

NMR Characterization of (ONBO) **21**:

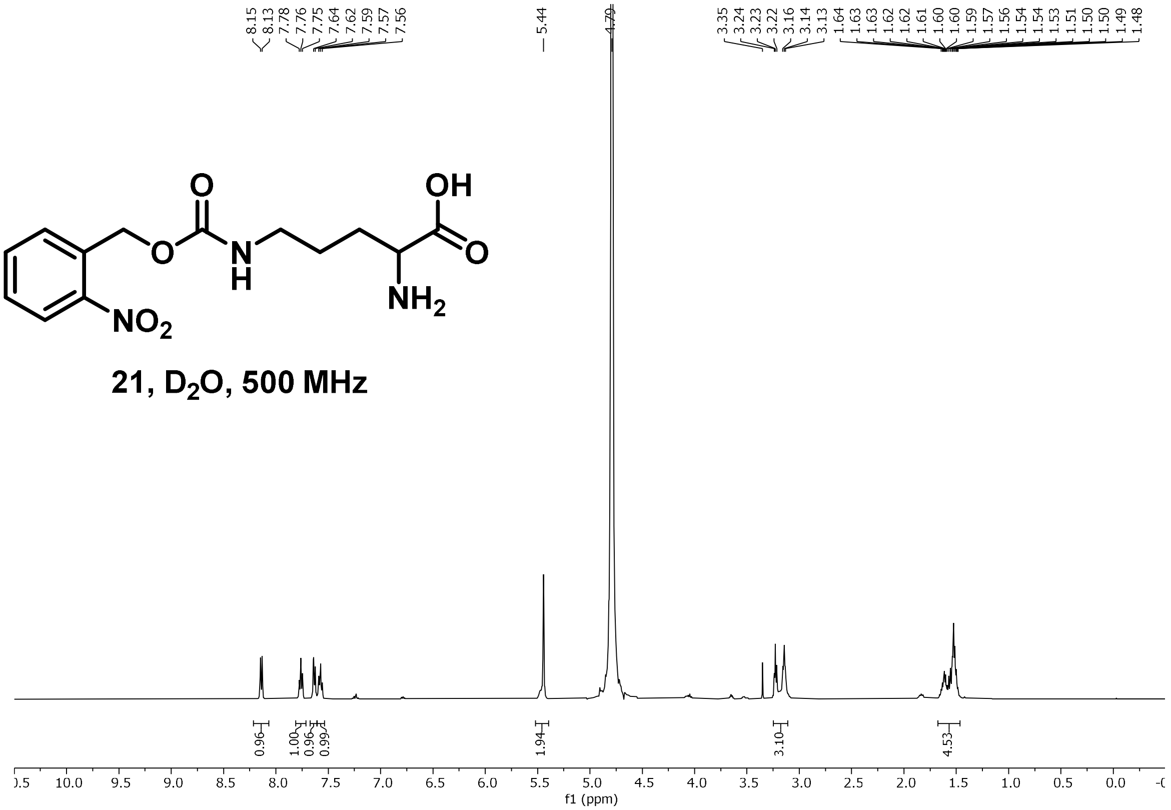


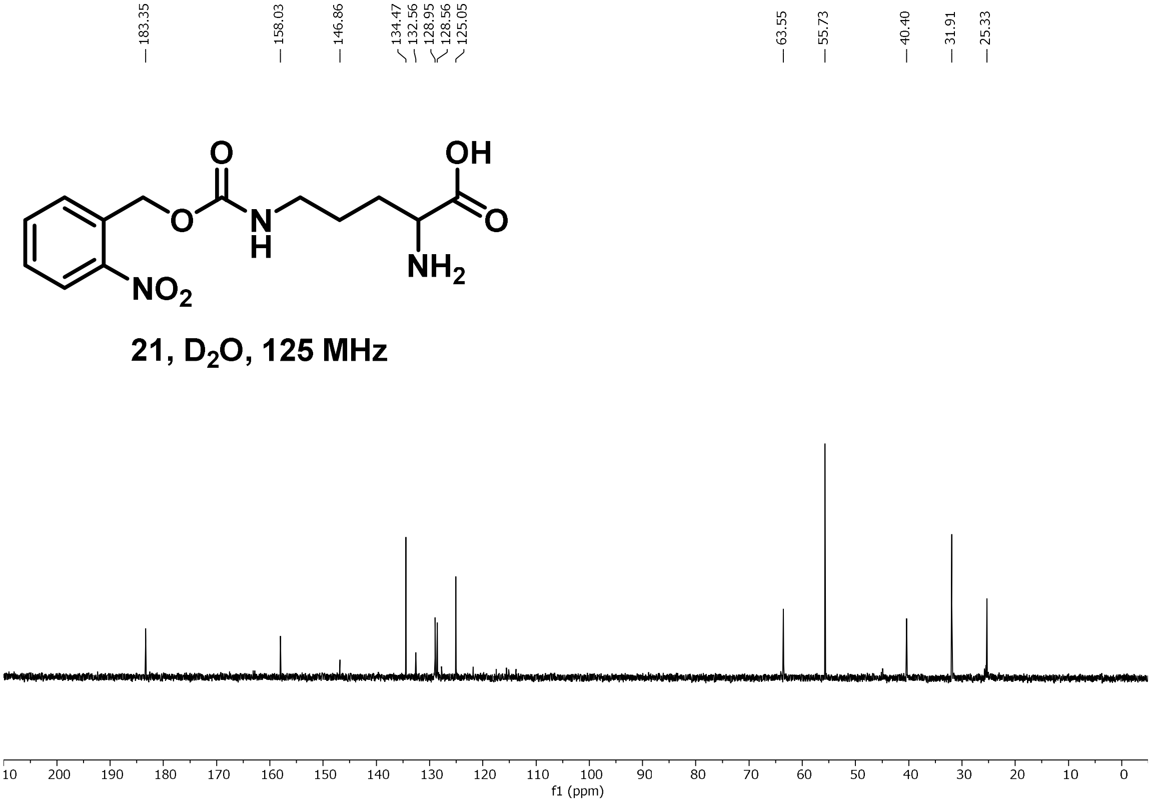

NMR Characterization of (Kacme) **3**:

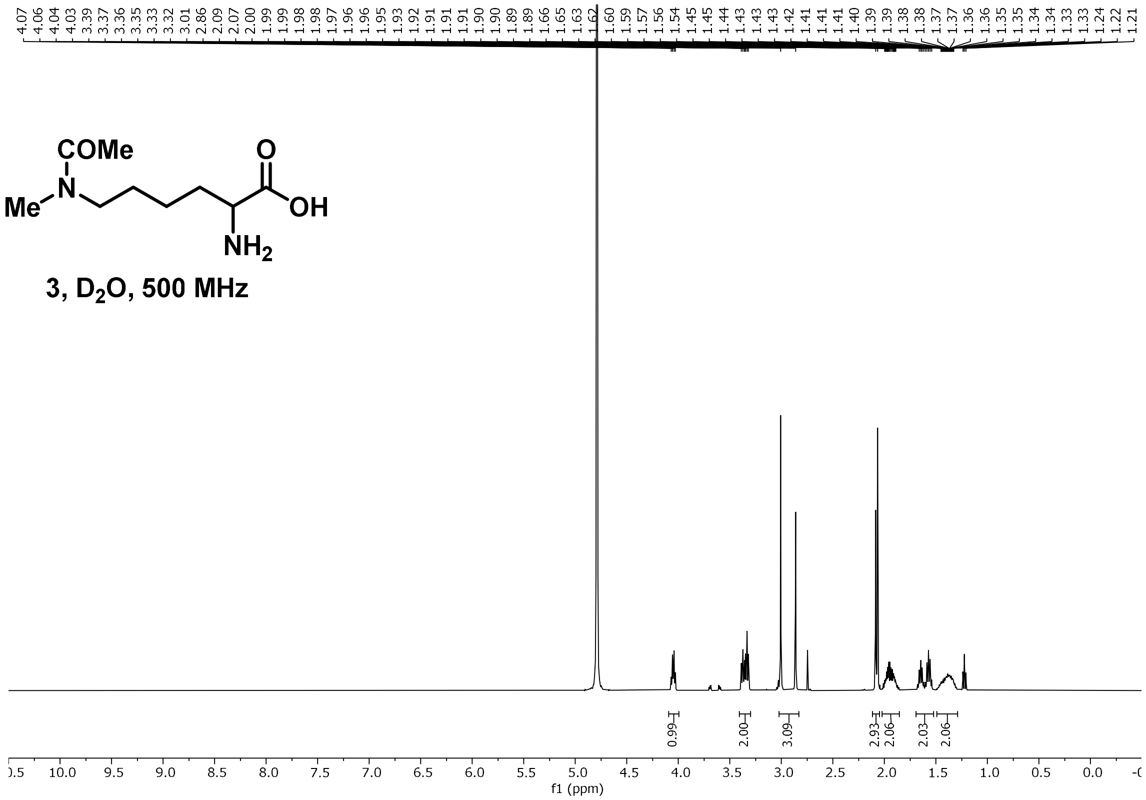


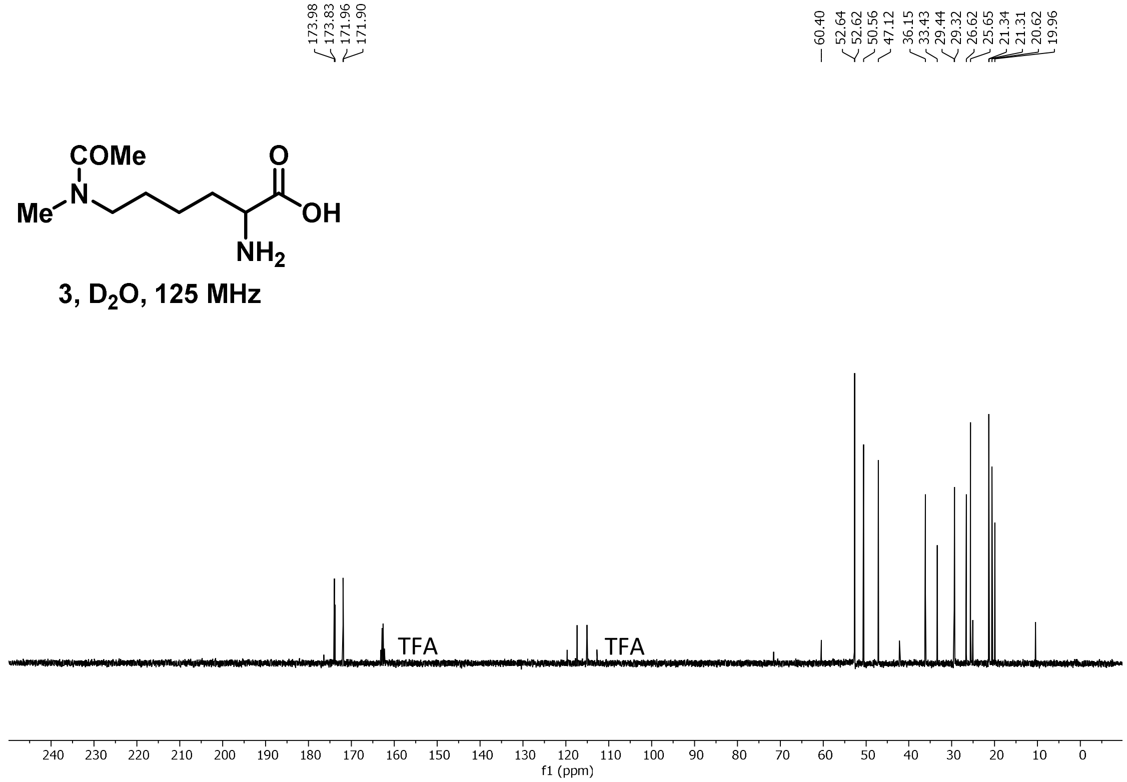

$^{1}H$ NMR of Kacme **13** at 80 °C

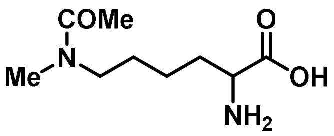

3, $D_2O$, 600 MHz, 353 K

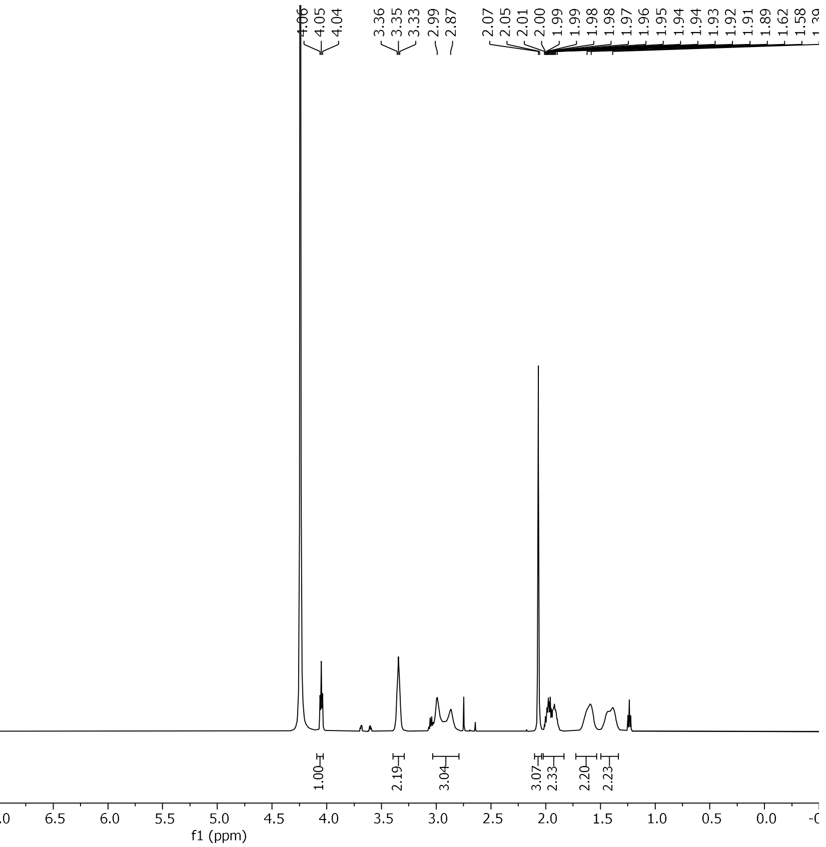
